# Simulation-guided sloppy DNA probe design for mismatch tolerant hybridization

**DOI:** 10.1101/2022.12.20.521289

**Authors:** Pallavi Bugga, Vishwaratn Asthana, Rebekah Drezek

## Abstract

The ability to both sensitively and specifically assess the sequence composition of a nucleic acid strand is an ever-growing field. Designing a detection scheme that can perform this function when the sequence of the target being detected deviates significantly from the canonical sequence however is difficult in part because probe/primer design is based on established Watson-Crick base-pairing rules. We present here a robust and tunable toehold-based exchange probe that can detect a sequence with a variable number of SNPs of unknown identity by inserting a series of controlled, sequential mismatches into the protector seal of the toehold probe, in an effort to make the protector seal “sloppy”. We show that the mismatch tolerant system follows predicted behavior closely even with targets containing up to four mismatches and thermodynamically deviating from the canonical sequence by up to 15 kcal/mole. The system also performs faithfully regardless of the global mismatch position on either the protector seal or target. Lastly, we demonstrate the generalizability of the approach by testing the increasingly sloppy protectors on HIV clinical samples and show that the system is capable of resolving multiple, iteratively mutated sequences derived from numerous HIV sub-populations with remarkable precision.

## Introduction

The detection of sequence-specific nucleic acids is a growing field not only in molecular biology but also the clinic.^1,2^ Increasing demand has been placed on nucleic acid-based detections probes to not only be sensitive and specific, but also robust and tunable. However, designing a probe *in silico* that meets these criteria when the sequence of the target being detected deviates significantly from the canonical sequence is incredibly difficult, in part because probe design is based on established Watson-Crick base-pairing rules.^3^ Tools like Next Generation Sequencing (NGS) can be used to determine the exact sequence identity of the region of interest, but such methods are often time intensive and costly.

While many clinically relevant single-nucleotide polymorphisms (SNPs) are well known, informing the design of specific detection probes, the identity of many SNPs are unknown *a priori*, thereby preventing the design of appropriate diagnostic probes.^4^ In certain circumstances, more than one SNP may be present in a region of interest, making reliable detection all the more difficult. This is especially true for hypermutable sequences including viral genomes, VDJ recombination regions, and trinucleotide repeats.^5–9^

To overcome this issue, we present a toehold-based probe system that can be intelligently designed *in silico* to detect a sequence with a variable number of mismatches or SNPs of unknown identity using thermodynamic cutoffs. Toehold probes are incredibly sensitive and specific nucleic acid-based detection tools that operate reliably under a wide range of salts and temperatures. In addition, relatively simple *in silico* design tools can be used to ensure toehold probes detect SNPs with high selectivity versus wildtype. However, because current iterations of the probe-protector system are designed to be ultra-specific, the toehold probes can only tolerate a known SNP either in the protector or target strand.^3,10^ When more than one SNP is present or the sequence is unknown, the system begins to deviate from its expected behavior. We show that introducing a series of controlled mismatches into the protector of the toehold probe, in an effort to make the protector “sloppy”, allows the protector to tolerate target sequences with an increasing number of mismatches/SNPs. By extension, the identities of these SNPs do not need to be known *a priori* to ensure robust detection.

While most nucleic acid-based detection probes are designed based on sequence homology to ensure specific binding, the sloppy protectors described herein rely predominantly on Gibbs free energy (ΔG°) cutoffs. Specifically, a target sequence with more mismatches than the sloppy protector will have a standard free energy of hybridization to the probe that is less favorable than the protector-probe duplex. As a result, the mismatched target cannot displace the protector and bind the probe. Conversely, a target with less mismatches and a more favorable ΔG° than the sloppy protector *can* displace the protector and bind to the probe. Despite using ΔG° cutoffs, the sloppy probe-protector system is still resilient to GC content, due to the requirement of target/protector sequence homology to probe.

Given that the average standard free energy penalty of a single-mismatch ranges from approximately 1.5 - 6.5 kcal/mole, and further that the hybridization efficiency of probe-protector displacement is sigmoidal in nature with respect to the standard free energy, with exponential-like qualities in the region between +5 to −5 kcal/mole (for or system), we can expect sharp cutoffs in hybridization yield for a set probe-protector complex with each target mismatch iteration.^10,11^ Accordingly, by using a set of increasingly sloppy protectors that span a sufficiently wide ΔG° range, the approximate ΔG° of the probetarget complex can be approximated and the identity of the target and its corresponding SNPs inferred. A schematic of the design is displayed in **Figure 1**.

**Fig. 1.**
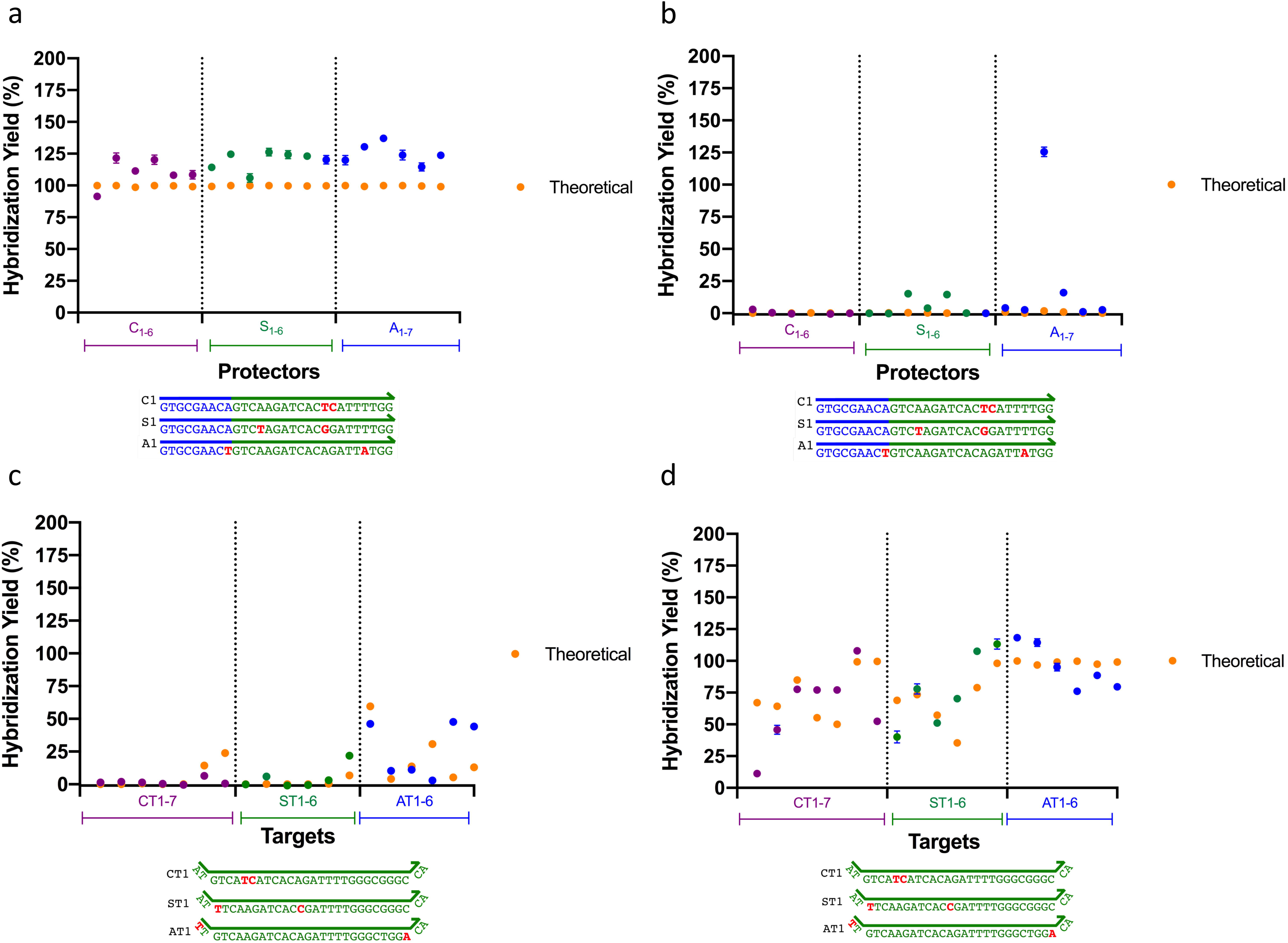
Schematic of Sloppy Protector Design Using X-Probes. **A)** Schematic of an X-Probe displacement reaction in the presence of a more complementary target. **B)** Theoretical hybridization yields for five increasingly sloppy protectors and targets. X-axis corresponds to the free energy difference between the probe-protector and probe-target, normalized to the free energy difference between the probe and C_3_. A target with more mismatches against the probe (relative to the protector) has a positive ΔΔG and will be less likely to displace the protector. Conversely, a target with less mismatches and a more favorable ΔG° than the sloppy protector can displace the protector and bind to the probe (ex. T_3_ to C_4-5_).

## Results

### Sloppy Protector Performance

To facilitate testing and reduce potential costs of using multiple sloppy protectors in parallel, X-probes developed previously were utilized.^12^ X-probes are conditionally fluorescent nucleic acid probes in which the two functionalized detection oligonucleotides (fluorescent [F] and quencher [Q]) are decoupled from the probe-protector-target complex. Consequently, the same F and Q species can be used with any combination of probe-protector detection pairs. In addition, the reaction mechanism of the X-Probe is similar to that of the toehold probe and well characterized.

Using NUPACK, five target [T_1-5_] and five protector (i.e. complement) sequences [C_1-5_] were designed, each with an increasing number of SNPs relative to the consensus probe [P] (T_1_ = zero mismatches ➜ T_5_ = four mismatches; C_1_ = zero mismatches ➜ C_5_ = four mismatches) (**Supplementary Table 1a and 1b**). The expected/theoretical yields of the various protector-target complexes is depicted in **Figure 2a**, and appears to correlate strongly with the experimental yields depicted in **Figure 2b**, indicating that current simulation methods can accurately approximate the thermodynamics of increasingly mismatched duplexes (adjusted-r^2^= 0.94, RMSE = 9.62). Experimental yields were calculated by normalizing each displacement against a positive control (the fluorescent strand only, or F_max_) with

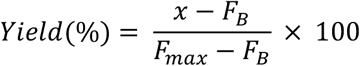

and where *x* corresponds to the end-point fluorescence of the reaction, and *F_B_* corresponds to the fluorescent background (the quenched X-probe without target added).

**Fig. 2.**
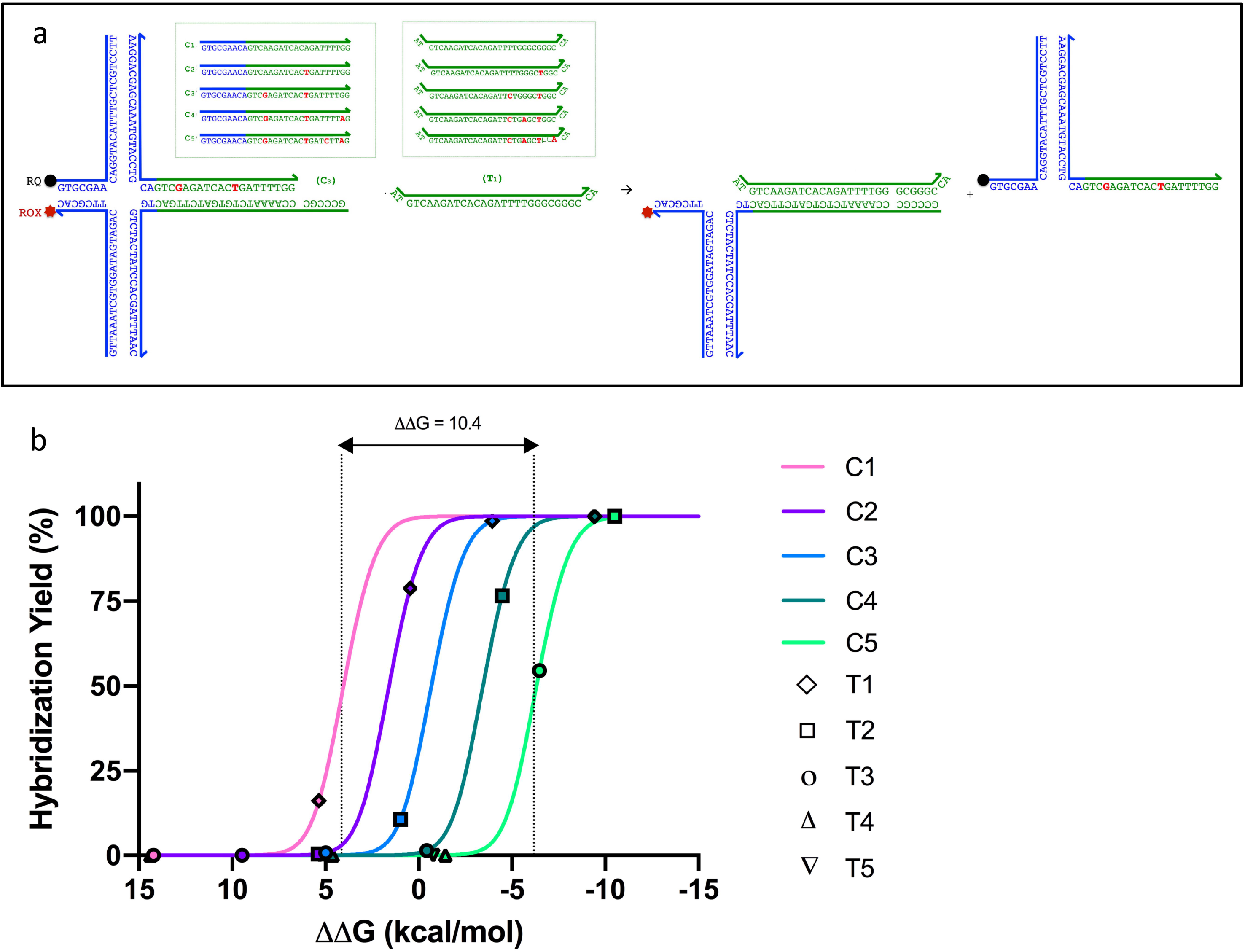
Experimental versus theoretical yields of increasingly sloppy mismatch tolerant hybridization probes. **A)** Expected/theoretical yields for five target (T_1-5_) and five protector (i.e. complement) sequences (C_1-5_), each with an increasing number of SNPs relative to the consensus probe (P) (T_1_ = zero mismatches ➜ T_5_ = four mismatches; C_1_ = zero mismatches ➜ C_5_ = four mismatches). Values are given in terms of percent yields. **B)** Experimental yields for five target (T_1-5_) and five protector (i.e. complement) sequences (C_1-5_), each with an increasing number of SNPs relative to the consensus probe (P) (T_1_ = zero mismatches ➜ T_5_ = four mismatches; C_1_ = zero mismatches ➜ C_5_ = four mismatches). Values are given in terms of percent yields. **C)** A linear plot of the data depicted in [B], demonstrates that hybridization of increasingly mismatched targets relative to progressively sloppy protectors follows strict ΔG° thresholds. Error bars represented the standard deviation of triplicate conditions.

**Figure 2c**, a linear plot of **Figure 2b**, demonstrates that hybridization and corresponding signal detection of increasingly mismatched targets relative to progressively sloppy protectors follows strict cutoffs that correspond to ΔG° thresholds. Further, kinetic traces of sloppy protector displacement indicate that the kinetics of the reaction are in agreement with previously characterized toehold probe reaction rate constants, and that increasing the number of mismatches in the target or protector strands has no appreciable effect on the rate of the strand displacement reaction, even when such mismatches occur on the target toehold, as is the case with targets T4 and T5 (**Supplementary Figures 1a-1e**).^10^ A table of all probe, protector, and target sequences and their respective ΔG° of hybridization can be found in **Supplementary Tables 1-2**.

### Impact of Global Mismatch Position

Next, to assess whether the global position of a mismatch, either on the protector or target, would disrupt predicted hybridization patterns and accordingly, the ability to accurately resolve ΔG° cutoffs, mismatches were inserted at set locations on the target and protector strand. Specifically, C_3_ protectors with two mismatches placed either immediately adjacent to one and another (C_1-6_), corresponding to zero bp of separation, seven bp apart (S_1-6_), or greater than 16 bp apart (A_1-7_) were tested against a T_1_ and T_3_ target (**Figures 3a, b, Supplementary Table 3**). Similarly, T_3_ targets with two mismatches spaced as described above were tested against a C_2_ and C_4_ protector (**Figures 3c, d, Supplementary Table 4**). Experimental versus simulation data confirms that the position of the mismatches on either the target or protector does not significantly alter predicted base-pairing thermodynamics and corresponding select displacement outside the specified ΔG° intervals, save for one outlier (A_4_ + T_3_). With the inclusion of said outlier, the adjusted r^2^ and RMSE equal 0.83 and 18.58, respectively. Upon elimination of the outlier, the adjusted r^2^ and RMSE become 0.89 and 15.00, respectively. Experimental yields for this experiment were calculated using the same method described above for Fig 1.

**Fig. 3.**
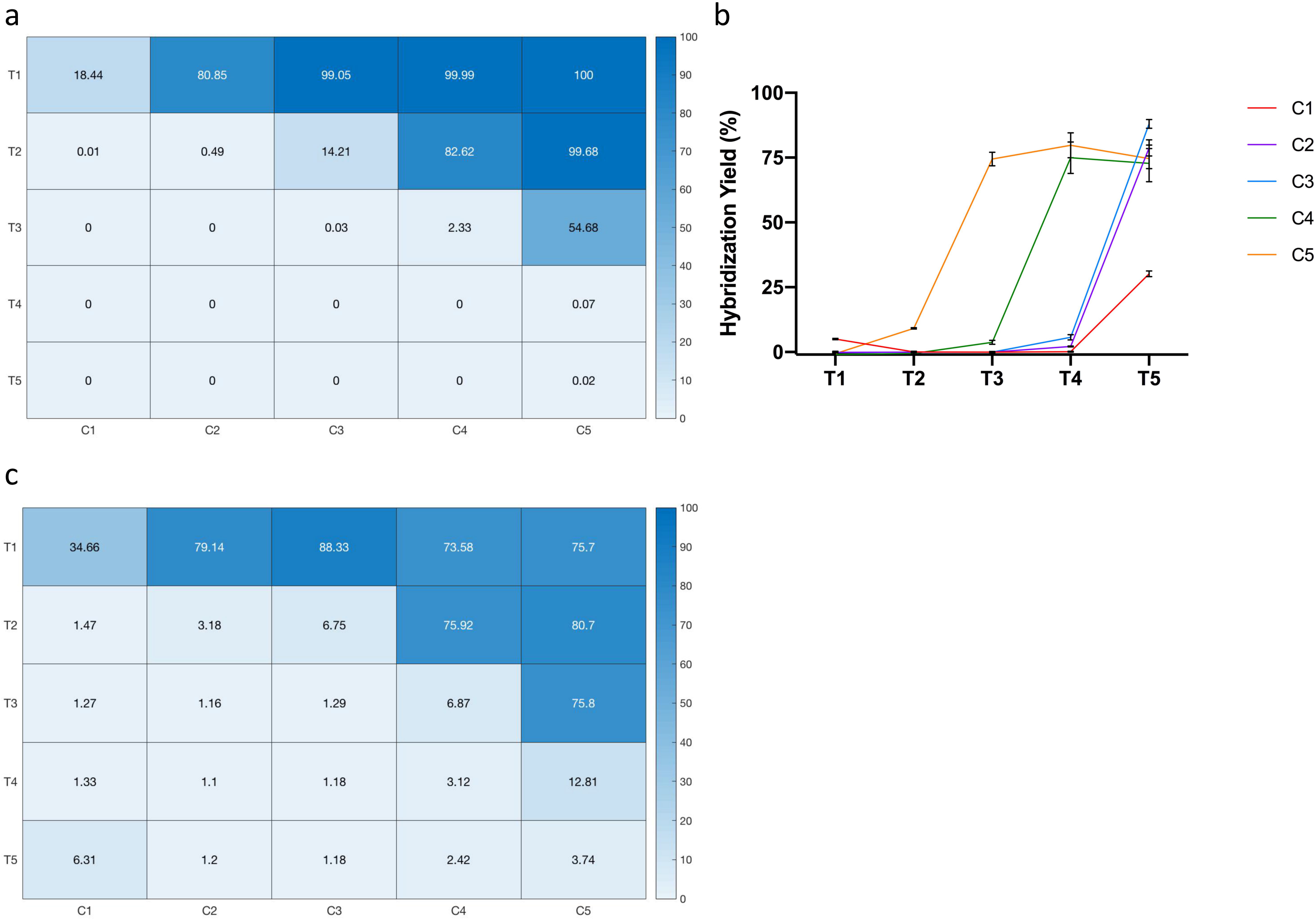
Assessing the impact of global mismatch position on sloppy protector performance. C_3_ protectors with two mismatches placed either immediately adjacent to one and another, corresponding to zero bp of separation (C_1-6_), seven bp apart (S_1-6_), or greater than 16 bp apart (A_1-7_) were tested against a **a)** T_1_ and **b)** T_3_ target. Similarly, T_3_ targets with two mismatches spaced as described above were tested against a **c)** C_2_ and **d)** C_4_ protector. Experimental yields versus simulation demonstrate the robustness of the mismatch tolerant hybridization system to global mismatch position, save for a few outliers. Error bars represented the standard deviation of triplicate conditions.

Overall, the representative 101 combinations of mismatched protectors and targets follow predicted behavior closely (r^2^=0.91, RMSE=13.46 without the above delineated outlier), validating the use of a set of increasingly sloppy protectors that span a sufficiently wide ΔG° range to approximate the ΔG° of a probe-target complex, which in turn can be used to infer the number and identity of a variable number of SNPs.

### Tests on Hypermutable Clinical Correlates

While the mismatch tolerant hybridization system thus far tolerated a single SNP-containing sequence that represents the only population in a sample, it is not clear how the system would behave in the presence of multiple sub-populations, a scenario frequently encountered in the setting of hypermutable genomes. For example, patients with longstanding HIV infection are known to harbor multiple subpopulations of virus, each typically containing an iterative number of mutations that were acquired as a result of the error-prone nature of HIV replication.^6,13^ To validate the robustness of the sloppy protectors in such a setting, we tested the system on HIV clinical sequences known to be highly mutable and generally difficult to characterize. HIV RNA was extracted from serum samples collected from HIV-infected patients and reverse transcribed into cDNA. Known hypermutable segments were sequenced using Next Generation Sequencing and a single region with significant heterogeneity in both number and type of SNPs was selected for testing using the sloppy protector system (**Supplementary Tables 5, 6**). Asymmetric PCR of the heterogeneous region was performed, yielding a 60 bp single-stranded target that was subsequently tested against newly designed X-probes (**Supplementary Table 7**). As shown in **Supplementary Figure 2a**, the differential displacement observed with increasingly sloppy protectors initially demonstrates moderate deviation from simulation (r^2^= 0.58, RMSE= 23.17). When a +5.5 kcal/mol penalty is imposed on the targets, however, the experimental results closely mirror simulation (r^2^= 0.99, RMSE= 4.006), as displayed in **Supplementary Figure 2b**. The rationale behind this penalty is explored further in the Discussion section. This result, however, demonstrates the generalizability of the system even for complex solutions.

## Discussion

We have demonstrated that a series of sloppy protectors that span a sufficiently wide ΔG° range can be used to approximate the ΔG° of a probe-target complex, which in turn can be used to infer the identity of a target and its corresponding SNPs. We start by showing that X-probes represent a relatively cheap and efficient method to couple multiple sloppy protectors with a single fluorophore-quencher strand complex obviating the need for repeated thermodynamic simulations of varying fluorophore-quencher paired sequences. Using NUPACK simulation software together with the X-probe design, five target and five protector sequences were designed, each with an increasing number of SNPs relative to the consensus probe. These SNPs were inserted at spaced intervals from one and another on both the protector and target. Further, where possible, the targets and protectors were designed with the following thermodynamic constraints in mind:

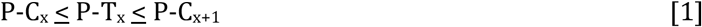

Given the sigmoidal nature of toehold displacement, where a marginal variation in ΔΔG° (defined as the ΔG° gap between [P-T] and [P-C]) significantly alters hybridization yields in and around null, while variations outside of this range contribute minimally, we can expect sharp cutoffs in displacement within each ΔG° interval delineated by equation [1]. This is especially true when the typical ΔG° penalty of a single mismatch varies anywhere between 1.5 - 6.5 kcal/mole, enabling the compartmentalization of individual mismatches within the intervals specified by each increasingly sloppy protector.^10,11^ Results of experiments testing T_1-5_ and C_1-5_ in Figure 2 confirm that hybridization of increasingly mismatched targets relative to progressively sloppy protectors follows strict cutoffs that correspond to ΔG° thresholds. Further, toehold displacement of these increasingly mismatched strands can be accurately predicted using current simulation methods.

Interestingly, experimental yields of displacement never reach 100%, even when predicted by theory, instead demonstrating an asymptotic maximum of roughly 80% fluorescence relative to the positive control. This trend has been previously observed, however, when evaluating the kinetics of strand displacement in X-probes by synthetic targets via fluorometer analysis, and is postulated to be a result of misaligned hybridization, or non-specific interactions between the target and probe strands.^11^ Further exploration of this effect may be warranted to fully elucidate the factors responsible for this reduced experimental maximum.

It should be noted that protectors C_2_ and C_3_ behave nearly identically in terms of their displacements to targets T_1-5_. For initial experiments, attempts were made to insert mismatches to not only follow the thermodynamic constraints depicted in equation [1] but also to be somewhat evenly spaced from one and another to minimize unintended deviations from predicted displacement. Because these design considerations are not mutually exclusive, the ΔG° gap between protectors C_2_ and C_3_ inadvertently ended up being smaller than the ΔG° gap between targets T_1_ and T_2_ leading to a slight deviation from equation [1] with P-T_1_ < P-C_2_ < P-C_3_ < P-T_2_ and a subsequent stagger (**Supplementary Table 1**). As a result, the thermodynamic cutoffs of protectors C_2_ and C_3_ overlap. The effects of this overlap however were accurately predicted via simulation.

Because the requirement to space mismatches evenly on the protector can interfere with the design of consistently spaced ΔG° gaps, and the position of multiple mismatches is not guaranteed to be evenly spaced on the target, we next assessed whether the global position of a mismatch, either on the protector or target, would disrupt predicted hybridization patterns and accordingly the selective displacement expected within each ΔG° interval. **Figures 3a** and **b** demonstrate that the presence of two mismatches stacked together, spaced 7 base pairs apart, or at the periphery of the protector strand (>16 base pairs apart) does not significantly alter select displacement of the C_3_ protector (C_3_ = two mismatches) relative to the T_1_ and T_3_ target, and similarly does not significantly alter select displacement of the T_3_ target (T_3_ = two mismatches) relative to the C_2_ and C_4_ protector relative to simulation. The bounding strands tested against the C_3_ protector and T_3_ target were chosen because their upper and lower thermodynamic affinities constrain the ΔG° range of the various mismatches.

It should be noted that of all the representative targets and protectors tested, a few outliers were evident where experimental displacement diverged from prediction. This is especially clear in **Figure 3b**, where the A_4_ variation of the C_3_ protector yields unanticipated displacement against T_3_ target. Secondary structure analysis of the A_4_ variation of the C_3_ protector demonstrates that the mismatches unique to these strands encourage hairpin formation at the 3’ end of the protector near the target toehold. As a result, the protector may have a propensity to self-dimerize as an inter-strand complex once strand displacement begins, incurring an additional thermodynamic penalty that permits the target to fully displace the protector. Interestingly, these protectors also demonstrate self-binding near the protector toehold region, perhaps hindering the ability of the protectors to re-bind the probe in equilibrium, leading to greater target-probe complexation stability. In addition, the A_4_ protector strand also contain mismatches immediately adjacent to the multi-loop junction, likely destabilizing the loop and allowing for more facile displacement of the protector from the probe. While these destabilizing mismatches are notably also present in protectors, A_1_, and A_6_, the combination of the effect of the mismatches with the above-described factors (that are specific to the A_4_ strand) may lead to the observed increase in protector displacement relative to theory seen with these cases. It should be noted that this effect is likely not immediately observed with A_4_ with T_1_, as the target is already thermodynamically favored to fully displace the A_4_ protector, and thus any added benefit to target-probe complexation does not significantly increase the expected yield. Other contributing factors that may lead to deviation from prediction include inaccuracies in NUPACK simulation, experimental error, and oligonucleotide synthesis errors, which may remain despite purification.^14,15^

The results obtained in **Figures 3a** and **3b** show yields of >100% for many of the reactions expected to show displacement (i.e. the mismatch protectors with T_1_ and the mismatch targets with C_4_). This yield is likely greater than experimental maximums (100%) due to relative photo-bleaching of the positive control—the fluorescent strand only— compared to the complexed X-probes. The complexed X-probes are stored in a quenched state, which may function to protect the fluorophores from photo-bleaching over time; the fluorescent positive control strands, on the other hand, remain unprotected during storage.

Altogether, the representative 101 combinations of mismatched protectors and targets follow predicted behavior closely demonstrating the robustness of the sloppy protector system and its potential as a previously unfeasible, finely tuned mismatch tolerant hybridization system. In addition, the set of protectors devised for the aforementioned experiments alone span a ΔG° range of 15 kcal/mol and together can accurately resolve variously mismatched targets within this interval with high fidelity. The range can theoretically be extended using an increasing number of longer and sloppier protectors with various options on how to space these strands. For example, successive sloppy protectors can be designed to have narrow ΔG° gaps to afford better resolution (A-T mismatch), or large ΔG° gaps to span a large detection range (G-C mismatch).

While the system was found to perform robustly in the presence of a single dominant sequence, containing a variable number and identity of SNPs, it was not yet clear how the system would behave in the presence of multiple sub-populations, each harboring sequences containing a unique SNP signature. The former scenario is typically encountered when SNPs are well conserved—such as in eukaryotes—in which the distribution of the unique SNP-containing sequence is generally expected to be present either at 100% (homozygous) or 50% (heterozygous) depending on the nature of that particular mutation.^16^ The latter situation, however, can be found in certain species with hypermutable sequences, such as viruses, in which several sequences with an iterative number of acquired SNPs of unknown identity can be typically expected to co-exist; these mutated genomes are often referred to as quasi-species, and have been investigated in the context of viral resistance to therapy.^6^ Based on sequencing data of the *pol* gene, taken from one of the three patient samples, three unique and related HIV subtypes or subpopulations were observed in this individual (at a prevalence of 69.15%, 25.76%, and 1.68% respectively), all of which likely began from an original strain that sequentially evolved and branched with chronic infection. The iterative nature of mutation acquisition has previously been demonstrated in the protease and reverse transcriptase domains of the *pol* gene of the HIV genome, as one of the driving forces behind resistance.^13^ Indeed our sequencing data seems to suggest that acquired mutations follow a sequential path, though there may be situations where this is not the case.

As shown in **Figure 4**, the sloppy protector design is capable of accurately resolving each unique HIV subtype as demonstrated by the incremental increase in signal corresponding to the relative distribution of each iteratively mutated sequence. Prior to adjustment, moderate deviation from simulation was noted. When a +5.5 kcal/mol penalty was imposed on the targets, however, the experimental results closely mirrored simulation. Though the nature of the +5.5 kcal/mol penalty has not been thoroughly investigated, based on initial simulations, we believe this penalty is a product of the complex secondary structure formed by the 60 bp single-stranded HIV region of interest, which must be disrupted to displace the protector and bind the probe. Truncating the length of the target being assessed has the potential to overcome this issue, as the degree of self-binding outside of the region of interest is reduced with decreasing strand length.

**Fig. 4.**
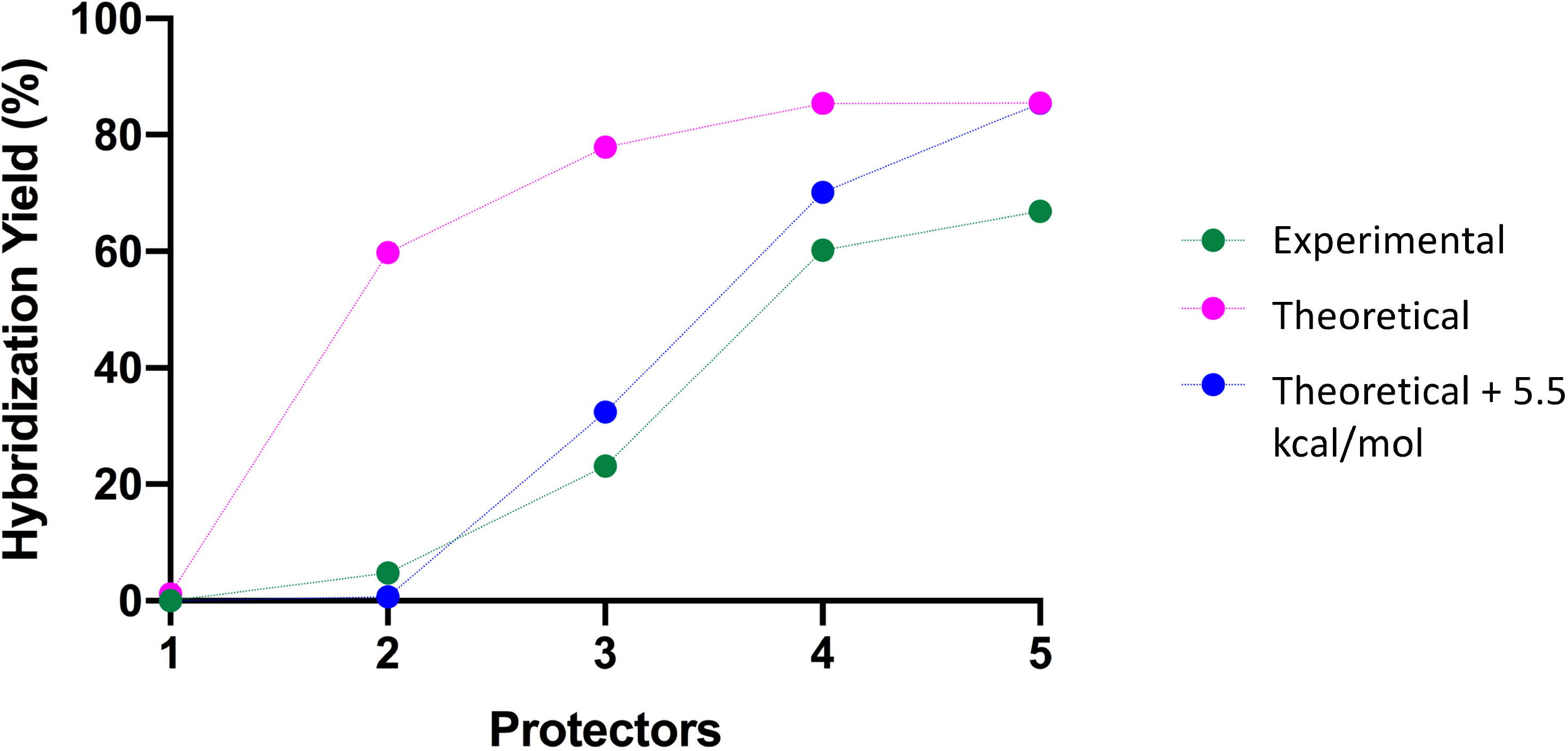
Evaluating Performance of Sloppy Protectors Against Hypermutable Clinical Correlates. Experimental yields are compared against theoretical yields, pre- and post-application of a 5.5 kcal/mol penalty. With the 5.5 kcal/mol penalty, experimental results correlate closely to theoretical estimates.

Without *a priori* knowledge of the sample's composition, it is theoretically possible to back-calculate the number of unique and increasingly mutated sequences in a sample, as well as the approximate ΔG° of each probe-sequence complex, which in turn can be used to infer the number and possible identity of a variable number of SNPs. In the case of HIV for example, patients who have developed resistant strains as a result of chronic, unchecked infection or treatment noncompliance, will either harbor a primary HIV genome that deviates from the endemic sequence or carry multiple HIV subpopulations as observed in this study - information that can help determine prognosis and guide therapy.

While still proof-of-concept, the number of potential applications of the mismatch tolerant hybridization system is still unrealized. Any hypermutable sequence or sequence with an uncharacterized set of SNPs can potentially be assayed using the sloppy protector system without the need to repeatedly test a sample using more costly approaches, such as sequencing. Further, unlike other hybridization-based assays—like microarrays—the system can tolerate mismatches in a finely tuned and precise manner without having to nonspecifically lower the hybridization temperature, adjust the salt concentration, or insert universal and/or more thermodynamically-favorable bases.^17–21^ Emerging technologies that utilize universal probes or primers also stand to benefit from such a design. As an example, the Universal Microbial Diagnostics system developed by previous members in our lab is a system that avails of the differential bindings of a set of universal, target-agnostic probes to various bacterial genomes as a means of identifying infectious bacterial species.^22^ While the initial scheme utilized sloppy molecular beacon probes, implementation of the mismatch-tolerant toehold probes described herein may improve the specificity of probe binding, and therefore increase the overall efficacy of the identification platform.

## Materials and Methods

### DNA Oligonucleotides

The DNA oligonucleotides used in this study were purchased from Integrated DNA Technologies (IDT, Coralville, IA). All purchased oligonucleotides were purified via standard desalting, and pre-diluted in pH 8.0 IDTE buffer at a concentration of 100 μM. The fluorescent and quencher strands were ordered with a Carboxy X-Rhodamine (ROX) fluorophore and Iowa Black RQ quencher 3’ and 5’ modification, respectively.

### X-Probe Annealing Protocol

X-probes were annealed at a ratio of 1:1.5:3:5 of fluorophore, probe, protector, and quencher respectively, using the following protocol: for a 100 μL reaction, added 10 μL 10 μM fluorescent strand, 15 μL 10 μM probe, 3 μL 100 μM protector, 5 μL 100 μM quencher strand, 50 μL 10x PBS, and 17 μL nuclease-free water. Strands were then run through the following thermocycling protocol on standard Eppendorf thermocyclers: incubation at 1) 95°C for 4 minutes, 2) 95°C for 1 minute, 3) ramp down by 1°C, 4) GOTO step 2 (repeat 75 ×), 5) 20°C for 1 minute.

### Fluorescent Quantification Studies

Fluorometer data from initial experiments testing sloppy protector performance was gathered on the Horiba Scientific Fluoromax-4 Spectrofluorometer. The optimal excitation and emission wavelengths for the ROX fluorophore were set at 582 nm and 600 nm, respectively. Slit sizes were fixed at 4 nm for both excitation and emission, and integration time was set to 10s (per cuvette) for each 60s time point. Temperature was maintained at 37°C.

To achieve a final concentration of 10 nM of X-Probe, 12 μL of the 1 μM X-Probe solution was mixed with 1200 μL of 5x PBS in a standard quartz cuvette. Prior to spiking the X-Probe solutions with target, five minutes of fluorescent background (f_B_) data was attained. Cuvettes were then removed from the apparatus and 24 μL of 1 μM target solution (20 nM final concentration) was added after which each cuvette was capped, inverted 20 times to mix, and placed back in the machine to allow data acquisition to continue. For acquisition of positive control fluorescence, 12 μL of 1 μM of fluorophore was added to 1200 μL of 5x PBS in a standard quartz cuvette, and allowed to incubate for 5 minutes in the apparatus before being measured.

Fluorescence data from experiments testing the effect of global mismatch position on sloppy protector performance were performed using a Bio-Rad CFX Connect Real-Time PCR Detection System (Bio-Rad, Hercules, CA). To set up the reaction, 10 μL of water, 10 μL of 1 μM target, 25 μL of 10x PBS + 0.2% Tween, and 5 μL of 1 μM X-Probe were combined and pipetted into a 96-well qPCR plate. Reactions were then allowed to incubate at 37°C for 1.5 hours. To process plates, end point ROX fluorescence was captured in the real-time PCR detection system using the following protocol: step 1) 37°C for one minute followed by fluorescent capture, step 2) GOTO step 1 × 5. RFU intensity readouts were then averaged over the five minutes. No background subtraction was performed.

### Preparation of HIV Clinical Samples

As an *in vitro* proof-of-concept, the efficacy of the sloppy toehold-probe system was evaluated in the context of resolving quasi-species or viral subpopulations present in clinical HIV. Clinical HIV samples were collected from patients prior to receiving ART treatment (IRB #: 18-05-1929). For all patients, HIV RNA counts in plasma were high (>750,000 copies/mL). RNA was extracted from plasma samples using the Qiagen QIAamp Viral RNA Mini Kit (Qiagen, Germantown, MD). RNA to cDNA conversion was performed using the SMART cDNA Library Construction Kit by Clontech Laboratories, Inc (Takara Bio, San Jose, CA). Converted cDNA products were next column purified and subsequently amplified using universal primers designed relative to an internal consensus sequence derived from analysis of 800 patient samples, and corroborated against the HIV HXB2 reference sequence (GenBank: K03455.1). Phusion PCR amplification was performed via the following protocol: 10 uL of 5x Phusion HF buffer, 0.4 uL of 25 mM dNTPs, 0.5 uL of Phusion Hot Start polymerase, 0.4 uL of 100 uM primer mastermix, 10 uL of template, and 28.7 uL of water were combined to form a 50 uL reaction (Phusion PCR reagents were purchased from New England Biolabs, Ipswich, MA). The thermocycling protocol proceeded first with an incubation at 98°C for 30 seconds, followed by 30 cycles at 98°C for 10 seconds, 63°C for 30 seconds, and 72°C for 3 minutes, and a final elongation step at 72°C for 5 minutes. Samples were then column purified once more to remove PCR by-products and residual enzyme/buffer.

### Hotspot Region Amplification of cDNA

cDNA samples were again purified via column purification using a Zymo Research DNA Clean and Concentrator-25 kit (Zymo, Irvine, CA). Select 200-300 bp regions were then amplified using specific primers designed against various hotspot regions found in the patient-sample derived consensus sequence (**Supplementary Table 5**), and a high-fidelity polymerase (Phusion) purchased from New England Biolabs (NEB, Ipswich, MA). 10 μL PCR reactions were set-up using 2 μL 5x HF Buffer (NEB), 0.5 μL 10 mM dNTPs (NEB), 0.5 μL 10 μM forward primer, 0.5 μL 10 μM reverse primer, 0.2 μL Phusion polymerase, and 4.5 μL water. Reactions were pipetted directly into 0.2 mL PCR tubes, vortexed, and centrifuged down. The thermocycling protocol was performed on a Bio-Rad T100 Thermal Cycler as follows: step 1) 98°C for 3 minutes; step 2) 98°C for 0.5 minutes; step 3) 63°C for 0.5 minutes; step 4) 72°C for 1 minute; Step 5) Repeat steps 2-4 45 times; Step 6) 72°C for 5 minutes; Step 7) 4°C Hold. After performing PCR amplification, samples were again purified via column purification using the Zymo Research DNA Clean and Concentrator-25 kit. Two rounds of PCR amplification were performed in total (using the above protocol). Amplicons were analyzed via Polyacrylamide Gel Electrophoresis (PAGE), quantified using a Qubit dsDNA fluorometer (Thermofisher Scientific, Waltham, MA), and sent to Genewiz for Sanger sequencing (Genewiz, South Plainfield, NJ). Based on the Sanger sequencing results, four regions—of the fourteen initially amplified (seven hotspot regions on two separate patient samples) were found to contain a sufficient distribution of mutations and were further processed for Next Generation Sequencing.

### Characterizing Clinical Isolates and Designing and Testing Corresponding Sloppy Probes

The four aforementioned PCR amplicons (analyzed via Sanger sequencing) were amplified once again using the same thermocycling protocol as used previously, and column purified. Post-amplification products were characterized via Bioanalyzer (Agilent, Santa Clara, CA) and quantified using the Qubit dsDNA Broad Range Assay kit. Samples were sequenced using the Next Generation Sequencing Amplicon-EZ service provided by Genewiz. Full sequencing results are available in the Supplementary section (**Supplementary Attachment 1 and 2**). A 30 bp segment in the Pair 1 amplicon was found demonstrating significant SNP diversity, for which X-probes were subsequently designed (see **Supplementary Table 6, 7**). Iteratively mutated sequence breakdowns for this segment were determined by evaluating all sequenced strands with a greater than 0.1% read rate over a 60 base pair region containing the 30 bp strand of interest. Asymmetric PCR was then performed on the sequenced sample to yield a 60-bp single-stranded “target” oligonucleotide (primers used are included in **Supplementary Table 8**). 0.5 uL template DNA (corresponding to a final concentration of roughly 1 nM) was added to 1 uL 1 uM reverse primer and 2 uL 10 uM forward primer (a 1:20 ratio), 10 uL 5x Phusion HF Buffer, 1 uL 10 mM dNTPs, 1 uL Phusion Polymerase, and 34.5 uL water to form a 50 uL reaction. This sample was then placed in a Bio-T100 Thermal Cycler to undergo the following thermo-cyling protocol: 98°C for 3 minutes, followed by 70 cycles at 98°C for 30 seconds, then 63°C for 1 minute, and 72°C for 1 minute, and then lastly, one final step of elongation at 72°C. PCR products were cleaned using the Zymo Clean and Concentrator-DNA 25 kit, and characterized using the Qubit dsDNA Broad Range assay kit, Qubit ssDNA assay kit, and a Nanodrop. To quantify displacement of the X-probes by the target strand, a reaction mixture consisting of 1200 uL 5x PBS + 0.1% Tween was combined with 6 uL of 100 nM of each X-Probe (for X-Probes 1-4) and 12 uL single-stranded target (synthesized via asymmetric PCR). Reactions were allowed to incubate for 1 hour at room temperature before being analyzed for 15 minutes in the Horiba Scientific Fluoromax-4 Spectrofluorometer as described previously (Horiba Instruments Incorporated, Irvine, CA).

### Standard Free Energy Calculation of Yields

For all experiments, the general strand displacement reaction can be approximated as:

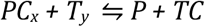

Where P represents the probe, C_x_ represents the complement strand or protector (C_1-5_), and T represents the target strand (T_1-5_). In this three-strand approximation, the region of the X-Probe immediately adjacent to the fluorophore/quencher label can be considered as contiguous with the probe and protector strands, and functions as the protector-toehold (8). The region of the probe and protector strands complementary to the fluorophore and quencher strands, respectively, are not considered when determining hybridization efficacy. For the three-strand approximation, *K_eq_* becomes

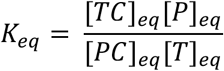

and can in turn be calculated from the difference between the free energies of the probe-complement complex and probe-target complex respectively (ΔΔG, where ΔΔG = ΔG_(PT)_ - ΔG _(PC))_ such that

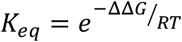

where *R* represents the universal gas constant and *T* represents temperature. ΔG_(PT)_ and ΔG_(PC)_ were determined using NUPACK simulation software (23), but can generally be calculated based on sequence composition using the algorithm described in (11).^11,23^ In our system, if we let *x* = [*TC*]_*eq*_, we find that [*PC*]_*eq*_ = [*PC*]_0_ – *x*, [*P*]_*eq*_ = [*P*]_0_ + *x*, and [*T*]_*eq*_ = [*P*]_0_ – *x*. Given our initial conditions ([*PC*]_0_ = 25 *nM*, [*P*]_0_ = 30 *nM* and [*P*]_0_ = 20 *nM*) we can rewrite *K_eq_* as

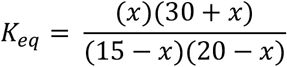

Solving for *x* allows for quantification of the hybridization yield *χ*, which can thusly be defined as the percent of probe-target relative to total probe concentration, or

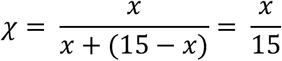

## Supporting information

Supplementary Attachment 1

## Acknowledgements

We would like to thank Drs. Roberto Arduino and Netanya Utay for providing the patient plasma samples (obtained under IRB protocol 18-05-1929). We would additionally like to thank Dr. Ruojia (Lucia) Wu for her assistance with processing, purifying, and reverse transcribing the patient plasma samples, as well as Dr. David Zhang for his guidance throughout this study. This research received no specific grant from any funding agency in the public, commercial, or not-for-profit sectors.

## Author Contributions

PB and VA designed all experiments; PB performed all experiments. PB and VA and RD analyzed all experimental data; PB and VA wrote and put together the manuscript; RD provided crucial intellectual input, and revised the manuscript. PB is the corresponding author for this submission; RD provided funding for this study. All authors read and approved the final manuscript.

## Data Availability Statement

The published article includes all datasets generated or analyzed during this study

