## Supplemental Tables and Figures for "Simulation-guided sloppy DNA probe design for mismatch tolerant hybridization"

### Supplementary Tables:

**Table S1a. Relevant X-Probe strands for Fig. 2 and 3.** The fluorophore (ROX) and quencher (RQ) are indicated in bold.

| Strand | Sequence |
|---|---|
| Probe | GCCCGCCCAAAATCTGTGATCTTGACTGGTCTACTATCCACGATTTAAC |
| Fluorescent Strand | GTTAAATCGTGGATAGTAGACTTCGCAC***ROX** |
| Quencher Strand | **RQ***GTGCGAACAGGTACATTTGCTCGTCCTT |

**Table S1b. ΔG of P+$C_{1-5}$ and P+$T_{1-5}$ for Fig.2.** Mismatches are indicated in bold.

| Sequence | dG (kcal/mol) |
|---|---|
| $C_1$: AAGGACGAGCAAATGTACCTGCAGTCAAGATCACAGATTTTGG | -37.8100 |
| $C_2$: AAGGACGAGCAAATGTACCTGCAGTCAAGATCAC**T**GATTTTGG | -35.3700 |
| $C_3$: AAGGACGAGCAAATGTACCTGCAGTC**G**AGATCAC**T**GATTTTGG | -33.0960 |
| $C_4$: AAGGACGAGCAAATGTACCTGCAGTC**G**AGATCAC**T**GATTTT**A**G | -30.3320 |
| $C_5$: AAGGACGAGCAAATGTACCTGCAGTC**G**AGATCAC**T**GAT**C**TT**A**G | -27.4606 |
| $T_1$: ATGTCAAGATCACAGATTTTGGGCGGGCCA | -37.2900 |
| $T_2$: ATGTCAAGATCACAGATTTTGGGC**T**GGCCA | -32.3500 |
| $T_3$: ATGTCAAGATCACAGATT**C**TGGGC**T**GGCCA | -28.3000 |
| $T_4$: ATGTCAAGATCACAGATT**C**TG**A**GC**T**GGCCA | -23.2600 |
| $T_5$: ATGTCAAGATCACAGATT**C**TG**A**GC**T**GG**A**CA | -22.5800 |

**Table S2. ΔΔG of P+$C_{1-5}$ and P+$T_{1-5}$ for Fig. 2**

| Target | C1(kcal/mol) | C2(kcal/mol) | C3(kcal/mol) | C4(kcal/mol) | C5(kcal/mol) |
|---|---|---|---|---|---|
| T1 | 0.52 | -1.92 | -4.19 | -6.96 | -9.83 |
| T2 | 5.46 | 3.02 | 0.75 | -2.02 | -4.89 |
| T3 | 9.51 | 7.07 | 4.80 | 2.03 | -0.84 |
| T4 | 14.55 | 12.11 | 9.84 | 7.07 | 4.20 |
| T5 | 15.23 | 12.79 | 10.52 | 7.75 | 4.88 |

**Table S3. Mismatch Protectors Used in Global Mismatch Position Experiments.** Mismatches are indicated in bold.

| Protector (Fig.3) | dG (kcal/mol) |
|---|---|
| C1: AAGGACGAGCAAATGTACCTGCAGTCAAGATCAC**TC**ATTTTGG | -31.98 |
| C2: AAGGACGAGCAAATGTACCTGCAGTCA**GA**ATCACAGATTTTGG | -31.88 |
| C3: AAGGACGAGCAAATGTACCTGCAGTCAAGATCACAGATT**CA**GG | -33.37 |
| C4: AAGGACGAGCAAATGTACCTGC**TT**TCAAGATCACAGATTTTGG | -31.93 |
| C5: AAGGACGAGCAAATGTACCTGCAGTCAAGAT**GT**CAGATTTTGG | -32.11 |
| C6: AAGGACGAGCAAATGTACCTGCAGTCAAGATCACAG**CA**TTTG | -33.03 |
| S1: AAGGACGAGCAAATGTACCTGCAGTC**T**AGATCAC**G**GATTTTGG | -32.93 |
| S2: AAGGACGAGCAAATGTACCTGCAG**A**CAAGATC**T**CAGATTTTGG | -31.61 |
| S3: AAGGACGAGCAAATGTACCTGC**T**GTCAAGA**G**CACAGCTTTTGG | -30.00 |
| S4: AAGGACGAGCAAATGTACCTGCAGTCAAGAT**G**ACAGATT**G**TGG | -31.45 |
| S5: AAGGACGAGCAAATGTACCTGCAGTCAAGA**A**CACAGAT**A**TTGG | -31.70 |
| S6: AAGGACGAGCAAATGTACCTGCAGTCAAG**C**TCACAGA**G**TTTGG | -32.36 |
| A1: AAGGACGAGCAAATGTACCTGC**T**GTCAAGATCACAGATT**A**TGG | -32.20 |
| A2: AAGGACGAGCAAATGTACCTGCA**T**TCAAGATCACAGATTTT**C**G | -30.88 |
| A3: AAGGACGAGCAAATGTACCTGCAG**C**CAAGATCACAGATTTTG**A** | -32.95 |

| | |
|---|---|
| A4: AAGGACGAGCAAATGTACCTG**T**AGTCAAGATCACAGAT**C**TTGG | -30.53 |
| A5: AAGGACGAGCAAATGTACCTGCA**C**TCAAGATCACAGATTT**G**GG | -30.90 |
| A6: AAGGACGAGCAAATGTACCTGC**T**GTCAAGATCACAGATTTT**A**G | -32.60 |
| A7: AAGGACGAGCAAATGTACCTGCAG**G**CAAGATCACAGATTTT**T**G | -33.03 |

**Table S4. Mismatch Targets Used in Global Mismatch Position Experiments.** Mismatches are indicated in bold.

| **Target (Fig.3)** | **dG (kcal/mol)** |
|---|---|
| CT1: ATGTCA**TC**ATCACAGATTTTGGGCGGGCCA | -31.63 |
| CT2: ATGTCAAGATC**TT**AGATTTTGGGCGGGCCA | -31.52 |
| CT3: ATGTCAAGATCACAG**TC**TTTGGGCGGGCCA | -32.49 |
| CT4: ATGTCAAGATCACAGATTT**CA**GGCGGGCCA | -31.19 |
| CT5: ATGTCAAGATCACAGATTTTGG**CG**GGGCCA | -31.01 |
| CT6: ATGTCAAGATCACAGATTTTGGGCGG**AG**CA | -34.63 |
| CT7: AT**AC**CAAGATCACAGATTTTGGGCGGGCCA | -35.10 |
| ST1: AT**T**TCAAGATCAC**C**GATTTTGGGCGGGCCA | -31.70 |
| ST2: ATGT**T**AAGATCACAG**G**TTTTGGGCGGGCCA | -31.89 |
| ST4: ATGTCA**T**GATCACAGATT**A**TGGGCGGGCCA | -31.26 |
| ST5: ATGTCAAGA**G**CACAGATTTT**A**GGCGGGCCA | -30.50 |
| ST6: ATGTCAAGATCACAG**G**TTTTGGGCGG**A**CCA | -32.15 |
| ST7: ATGTCAAGATCACAGATT**A**TGGGCGGGCC**T** | -34.06 |
| AT1: **T**TGTCAAGATCACAGATTTTGGGCGGG**A**CA | -36.38 |
| AT2: A**C**GTCAAGATCACAGATTTTGGGCGG**C**CCA | -33.70 |
| AT3: A**A**TTCAAGATCACAGATTTTGGGCGGG**A**CA | -34.59 |
| AT4: ATG**C**CAAGATCACAGATTTTGGGCGGGC**T**A | -35.37 |
| AT5: ATGT**T**AAGATCACAGATTTTGGGCGGGC**G**A | -33.88 |
| AT6: ATGTC**T**AGATCACAGATTTTGGGCGGGC**T**A | -34.55 |

*mismatches are bolded*

**Table S5. Hotspot Amplification Primers for Clinical HIV Samples**

| **Forward Primer** | **Corresponding Reverse Primer** |
|---|---|
| AATACATACTGACAATGGCAG | CTTTCCCCTGCACTGTA |
| TCTCTGGAACAGATTTGGAAT | CTTAAACCTACCAAGCCTCC |
| GGGAATCAAGCAGGAATT | CTTTCCCCTGCACTGTA |
| TAGAAGCAGAAGTTATTCCAGC | GATGAATACTGCCATTTGTACTG |
| TTGTACACATTTAGAAGGAAAAGT | TAGGGAATTCCAAATTCCTG |
| AGTGAAATTATGGTACCAGTTAGAG | GCATATTGTGAGTCTGTTACTATG |
| ATGATGTAAAACAATTAACAGAGG | GTAACATATCCTGCTTTTCCTAATT |

**Table S6. Percent Distribution of Unique HIV Viral Quasi-Species in a Human HIV Patient Sample.** Reads with a percent prevalence less than 0.1% were eliminated as they were below the reported Illumina sequencing error rate. Sequence similarities within the 60 bp region of interest were evaluated and re-tabulated to produce a condensed table of reads. New percent prevalence values were then calculated against said table.

| **% Prevalence** | **Sequence** |
|---|---|
| 69.15% | GTGGGCAGGGATTAAGCAGGAATTTGGCAT |
| 25.76% | GTGGGCAGGGATTAA**T**CAGGAATTTGGCAT |
| 1.68% | GTGGGCAGGG**G**TTAA**T**CAGGAATTTGGCAT |

**Table S7. X-probe strands for Elucidation of HIV Viral Quasi-Species in Human HIV Patient Sample.** Mismatches are indicated in bold.

| Strand | Sequence |
|---|---|
| Probe | GCCAAATTCCTGCTTAATCCCTGCCCTGGTCTACTATCCACGATTTAAC |
| Fluorescent Strand | GTTAAATCGTGGATAGTAGACTTCGCAC***ROX*** |
| Quencher Strand | **RQ***GTGCGAACAGGTACATTTGCTCGTCCTT |
| Protector 1 | AAGGACGAGCAAATGTACCTGCAGGGCAGGGATTAAGCAGGAA |
| Protector 2 | AAGGACGAGCAAATGTACCTGCAGGGCAG**A**GATTAAGCAGGAA |
| Protector 3 | AAGGACGAGCAAATGTACCTGCAGGGCAG**A**GATT**C**AGCAGGAA |
| Protector 4 | AAGGACGAGCAAATGTACCTGCAGGGCAG**A**GATT**C**AGCA**T**GAA |
| Protector 5 | AAGGACGAGCAAATGTACCTGCAG**T**GCAG**A**GATT**C**AGCA**T**GAA |

**Table S8. PCR primers for Asymmetric Amplification of HIV Hotspot Target**

| Primers | Amplicon |
|---|---|
| F: CAGTTAAGGCCGCCTGTT | CAGTTAAGGCCGCCTGTTGGTGGGCAGGGATTAAGCAGGAATTT GGCATTCCCTACAATC |
| R: GATTGTAGGGAATGCCAAA | |

**Table S9**. **Next-generation sequencing data for clinical HIV patient samples.** Please see attached Excel file.

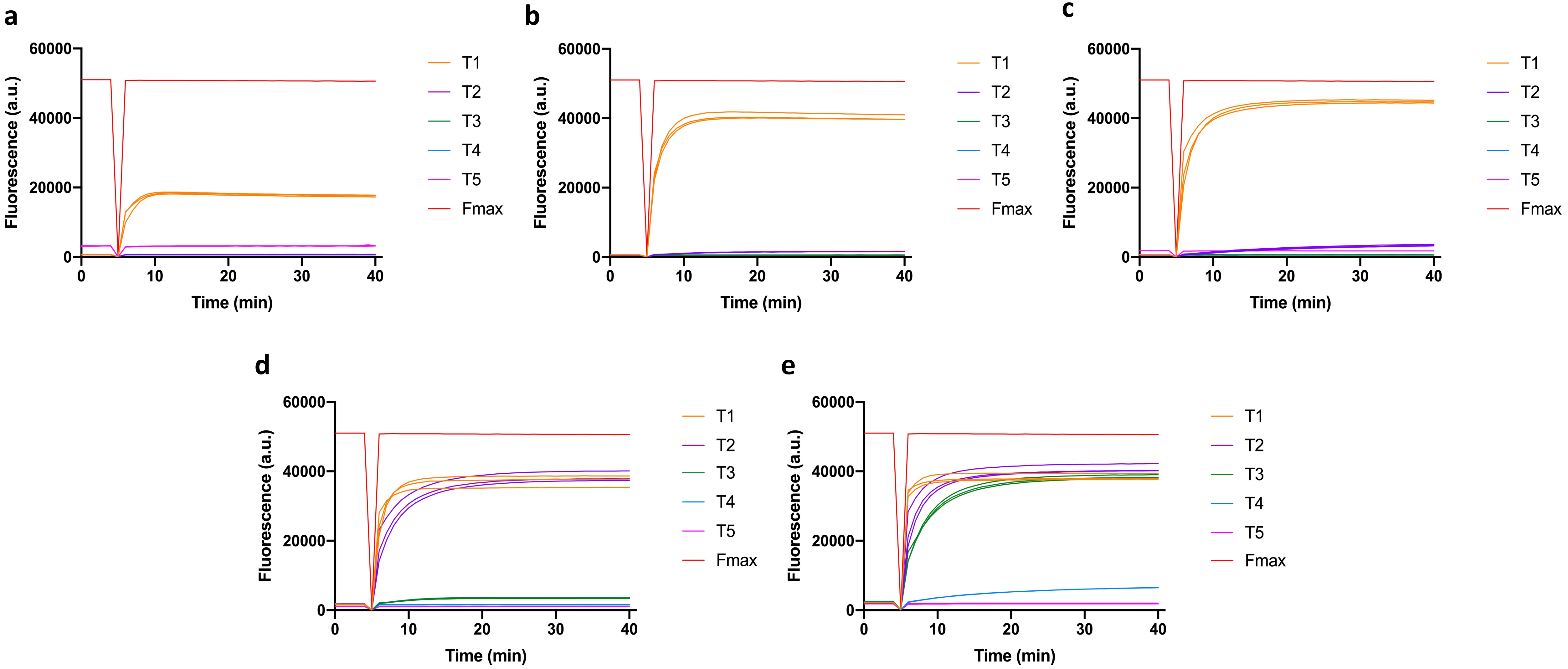


**Figure S1**. **Kinetic traces of "sloppy" X-probe strand - displacement reactions**. Traces correspond to reactions of protector $C_1$ with targets **(a)** $T_1$, **(b)** $T_2$, **(c)** $T_3$, **(d)** $T_4$, and **(e)** $T_5$. Kinetic traces demonstrate that "sloppy" strand displacement reaction kinetics are in agreement with previously characterized toehold probe reaction rate constants. Further, the number of mismatches on the target or protector strand does not affect the rate of strand displacement, even when the mismatches are present on the target toehold, as is the case with $T_4$ and $T_5$.

a

b

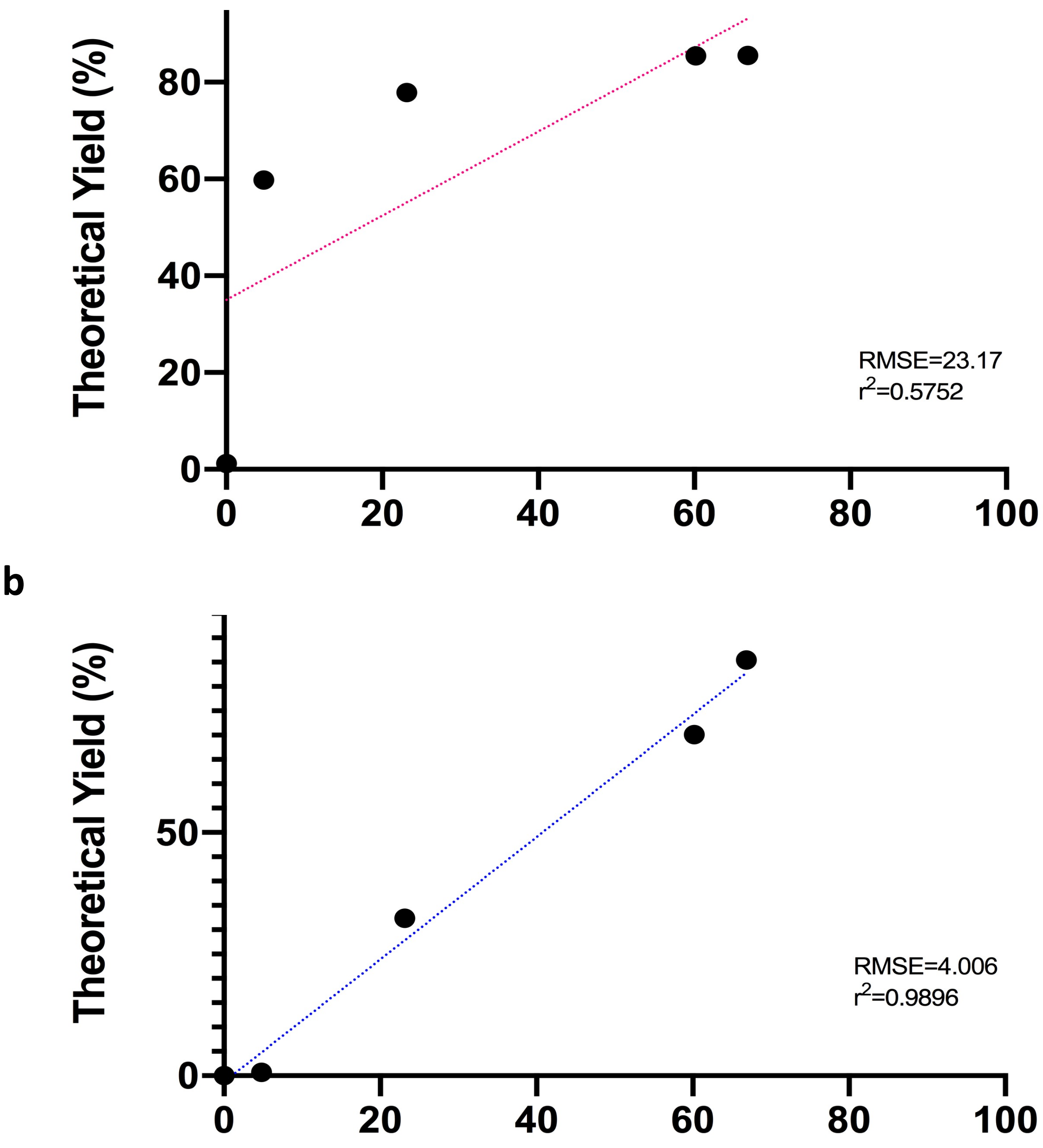


**Figure S2. Further Evaluation of the Performance of Sloppy Protectors Against Hypermutable Clinical Correlates a)** Linear regression of experimental versus theoretical yields, in the absence of the 5.5 kcal/mol penalty. X-axis represents experimental yield (%). **b)** Linear regression of experimental versus theoretical yields with the addition of the 5.5 kcal/mol penalty. Comparison of (a) and (b) demonstrates a significant improvement in correlation, based on Pearson coefficients, after the addition of the penalty.
