## Supplementary Attachment 1 for "Simulation-guided sloppy DNA probe design for mismatch tolerant hybridization"

**Supplementary Attachment 1. Next-generation sequencing results for Pair 1 Amplicon**

TargetSequence

TAGAAGCAGAAGTTATTCCAGCAGAAACAGGACAGGAAACAGCATACTTTATCTTAAAACTAGCAGGAAGATGGCCAGTAAAAACAATACATACAGACAATGGCAGCAATTTTACCAGTAATACAGTTAAGGCCGCCTGTTGGTGGGCAGGGATTAAGCAGGAATTTGGCATTCCCTACAATCCCCAAAGTCAGGAGTAGTAGAATCTATGAATAAAGAATTAAAGAAAATTATAGGACAGGTAGAGAACAAGCTGAACATCTTAAGACAGCAGTACAAATGGCAGTATTCATC

TAGAAGCAGAAGTTATTCCAGCAGAAACAGGACAGGAAACAGCATACTTTATCTTAAAACTAGCAGGAAGATGGCCAGTAAAAACAATACACACAGACAATGGCAGCAATTTTACCAGTAATACAGTTAAGGCCGCCTGTTGGTGGGCAGGGATTAATCAGGAATTTGGCATTCCCTACAATCCCCAAAGTCAGGAGTAGTAGAATCTATGAATAAAGAATTAAAGAAAATTATAGGACAGGTAGAGAACAAGCTGAACATCTTAAGACAGCAGTACAAATGGCAGTATTCATC

TAGAAGCAGAAGTTATTCCAGCAGAAACAGGACAGGAAACAGCATACTTTATCTTAAAACTAGCAGGAAGATGGCCAGTAAAAACAATACACACAGACAATGGCAGCAATTTTACCAGTAATACAGTGAAGGCCGCCTGTTGGTGGGCAGGGATTAAGCAGGAATTTGGCATTCCCTACAATCCCCAAAGTCAGGAGTAGTAGAATCTATGAATAAAGAATTAAAGAAAATTATAGGACAGGTAGAGAACAAGCTGAACATCTTAAGACAGCAGTACAAATGGCAGTATTCATC

TAGAAGCAGAAGTTATTCCAGCAGAAACAGGACAGGAAACAGCATACTTTATCTTAAAACTAGCAGGAAGATGGCCAGTAAAAACAATACACACAGACAATGGCAGCAATTTTACCAGCAATACAGTTAAGGCCGCCTGTTGGTGGGCAGGGATTAAGCAGGAATTTGGCATTCCCTACAATCCCCAAAGTCAGGAGTAGTAGAATCTATGAATAAAGAATTAAAGAAAATTATAGGACAGGTAGAGAACAAGCTGAACATCTTAAGACAGCAGTACAAATGGCAGTATTCATC

TAGAAGCAGAAGTTATTCCAGCAGAAACAGGACAGGAAACAGCATACTTTATCTTAAAACTAGCAGGAAGATGGCCAGTAAAAACAATACACACAGACAATGGCAGCAATTTTACCAGTAATACAGTTAAGGCCGCCTGTTGGTGGGCAGGGGTTAATCAGGAATTTGGCATTCCCTACAATCCCCAAAGTCAGGAGTAGTAGAATCTATGAATAAAGAATTAAAGAAAATTATAGGACAGGTAGAGAACAAGCTGAACATCTTAAGACAGCAGTACAAATGGCAGTATTCATC

TAGAAGCAGAAGTTATTCCAGCAGAAACAGGACAGGAAACAGCATACTTTATCTTAAAACTAGCAGGAAGATGGCCAGTAAAAACAATACATACAGACAATGGCAGCAATTTTACCAGTAATACAGTTAAGGCCGCCTGTTGGTGGGCAGGGATTAAGCAGGAATTTGGCATTCCCTACAATCCCCAAAGTCAGGAGTAGTAGAATCTATGAATAAAGAATTAAAGAAAATTATAGGACAGGTAGAGAACAAGCTGAACATCTTAAGACAGCAGTACAAATGGCAGTATTCATC

TAGAAGCAGAAGTTATTCCAGCAGAAACAGGACAGGAAACAGCATACTTTATCTTAAAACTAGCAGGAAGATGGCCAGTAAAAACAATACACACAGACAATGGCAGCAATTTTACCAGTAATACAGTTAAGGCCGCCTGTTGGTGGGCAGGGATTAAGCAGGAATTTGGCATTCCCTACAATCCCCAAAGTCAGGAGTAGTAGAATCTATGAATAAAGAATTAAAGAAAATTATAGGACAGGTAGAGAACAAGCTGAACATCTTAAGACAGCAGTACAAATGGCAGTATTCATC

TAGAAGCAGAAGTTATTCCAGCAGAAACAGGACAGGAAACAGCATACTTTATCTTAAAACTAGCAGGAAGATGGCCAGTAAAAACAATACACACAGACAATGGCGGCAATTTTACCAGTAATACAGTTAAGGCCGCCTGTTGGTGGGCAGGGATTAAGCAGGAATTTGGCATTCCCTACAATCCCCAAAGTCAGGAGTAGTAGAATCTATGAATAAAGAATTAAAGAAAATTATAGGACAGGTAGAGAACAAGCTGAACATCTTAAGACAGCAGTACAAATGGCAGTATTCATC

TAGAAGCAGAAGTTATTCCAGCAGAAACAGGACAGGAAACAGCATACTTTATCTTAAAACTAGCAGGAAGATGGCCAGTAAAAACAATACACACAGACAATGGCAGCAATTTTACCAGTAATACAGTTAAGGCCGCCTGTTGGTGGGCAGGGATTAAACAGGAATTTGGCATTCCCTACAATCCCCAAAGTCAGGAGTAGTAGAATCTATGAATAAAGAATTAAAGAAAATTATAGGACAGGTAGAGAACAAGCTGAACATCTTAAGACAGCAGTACAAATGGCAGTATTCATC

TAGAAGCAGAAGTTATTCCAGCAGAAACAGGACAGGAAACAGCATACTTTATCTTAAAACTAGCAGGAAGATGGCCAGTAAAAACAATACACACAGACAATGGCAGCAATTTTACCAGTAATACAGTTAAGGCCGCCTGTTGGTGGGCAGGGATTAATCAGGAATTTGGCATTCCCTACAATCCCCAAAGTCAGGAGTAGTAGAATCTATGAATAAAGAATTAAAGAAAATTATAGGACAGGTAGAGAACAAGCTGAACATCTTAAGACAGCAGTACAAATGGCAGTATTCATC

TAGAAGCAGAAGTTATTCCAGCAGAAACAGGACAGGAAACAGCATACTTTATCTTAAAACTAGCAGGAAGATGGCCAGTAAAAACAATACACACAGACAATGGCAGCAATTTTACCAGTAATACAGTTAAGGCTGCCTGTTGGTGGGCAGGGATTAAGCAGGAATTTGGCATTCCCTACAATCCCCAAAGTCAGGAGTAGTAGAATCTATGAATAAAGAATTAAAGAAAATTATAGGACAGGTAGAGAACAAGCTGAACATCTTAAGACAGCAGTACAAATGGCAGTATTCATC

TAGAAGCAGAAGTTATTCCAGCAGAAACAGGACAAGAAACAGCATACTTTATCTTAAAACTAGCAGGAAGATGGCCAGTAAAAACAATACACACAGACAATGGCAGCAATTTTACCAGTAATACAGTTAAGGCCGCCTGTTGGTGGGCAGGGATTAAGCAGGAATTTGGCATTCCCTACAATCCCCAAAGTCAGGAGTAGTAGAATCTATGAATAAAGAATTAAAGAAAATTATAGGACAGGTAGAGAACAAGCTGAACATCTTAAGACAGCAGTACAAATGGCAGTATTCATC

TAGAAGCAGAAGTTATTCCAGCAGAAACAGGACAGGAAACAGCATACTTTATCTTAAAACTAGCAGGAAGATGGCCAGTAAAAACAATACACACAGACAATGGCAGCAATTTTACCAGTAATACAGTTAAGGCCGCCTGTTGGTGGGCAGGGATTAAGCAGGAATTTGGTATTCCCTACAATCCCCAAAGTCAGGAGTAGTAGAATCTATGAATAAAGAATTAAAGAAAATTATAGGACAGGTAGAGAACAAGCTGAACATCTTAAGACAGCAGTACAAATGGCAGTATTCATC

TAGAAGCAGAAGTTATTCCAGCAGAAACAGGACAGGAAACAGCATACTTTATCTTAAAACTAGCAGGAAGATGGCCAGTAAAAACAATACACACAGACAATGGCAGCAATTTTACCAGTAATACAGTTAAGGCCGCCTGTTGGTGGGCAGGGATTAAGCAGGAATTTGGCATTCCCTACAATCCCCAAAGTCAGGAGTAGTAGAATCTATGAATAAAGAATTAAAGAAAATTATAGGACAGGTAGAGAACAAGCTGAACATCTTAAGACAGCAGTACAAATGGCAGTATTCATC

TAGAAGCAGAAGTTATTCCAGCAGAAACAGGACAGGAAACAGCATACTTTATCTTAAAACTAGCAGGAAGATGGCCAGTAAAAACAATACACACAGACAATGGCAGTAATTTTACCAGTAATACAGTTAAGGCCGCCTGTTGGTGGGCAGGGATTAAGCAGGAATTTGGCATTCCCTACAATCCCCAAAGTCAGGAGTAGTAGAATCTATGAATAAAGAATTAAAGAAAATTATAGGACAGGTAGAGAACAAGCTGAACATCTTAAGACAGCAGTACAAATGGCAGTATTCATC

TAGAAGCAGAAGTTATTCCAGCAGAAACAGGACAGGAAACAGCATACTTTATCTTAAAACTAGCAGGAAGATGGCCAGTAAAAACAATACACACAGACAATGGCAGCAATTTTACCAGTAATACAGTTAAGGCCGCCTGTTGGTGGGCAGGGATTAAGCAGGAATTTGGCATTCCCTACAATCCCCAAAGTCAGGAGTAGTAGAATCTATGAATAAAGAATTAAAGAAAATTATAGGACAGGTAAAAGAACAAGCTGAACATCTTAAGACAGCAGTACAAATGGCAGTATTCATC

TAGAAGCAGAAGTTATTCCAGCAGAAACAGGACAGGAAACAGCATACTTTATCTTAAAACTAGCAGGAAGATGGCCAGTAAAAACAACACACACAGACAATGGCAGCAATTTTACCAGTAATACAGTTAAGGCCGCCTGTTGGTGGGCAGGGATTAAGCAGGAATTTGGCATTCCCTACAATCCCCAAAGTCAGGAGTAGTAGAATCTATGAATAAAGAATTAAAGAAAATTATAGGACAGGTAGAGAACAAGCTGAACATCTTAAGACAGCAGTACAAATGGCAGTATTCATC

TAGAAGCAGAAGTTATTCCAGCAGAAACAGGACAGGAAACAGCATACTTTATCTTAAAACTAGCAGGAAGATGGCCAGTAAAAACAATACACACAGACAATGGCGGCAATTTTACCAGTAATACAGTTAAGGCCGCCTGTTGGTGGGCAGGGATTAATCAGGAATTTGGCATTCCCTACAATCCCCAAAGTCAGGAGTAGTAGAATCTATGAATAAAGAATTAAAGAAAATTATAGGACAGGTAGAGAACAAGCTGAACATCTTAAGACAGCAGTACAAATGGCAGTATTCATC

TAGAAGCAGAAGTTATTCCAGCAGAAACAGGACAGGAAACAGCATACTTTATCTTAAAACTAGCAGGAAGATGGCCAGTAAAAACAATACACACAGACAATGGCAGCAATTTTACCAGTAATACAGTTAAGGCCGCCTGTTGGTGGGCAGGGATTAAGCAGGAGTTTGGCATTCCCTACAATCCCCAAAGTCAGGAGTAGTAGAATCTATGAATAAAGAATTAAAGAAAATTATAGGACAGGTAGAGAACAAGCTGAACATCTTAAGACAGCAGTACAAATGGCAGTATTCATC

TAGAAGCAGAAGTTATTCCAGCAGAAACAGGACAGGAAACAGCATACTTTATCTTAAAATTAGCAGGAAGATGGCCAGTAAAAACAATACACACAGACAATGGCAGCAATTTTACCAGTAATACAGTTAAGGCCGCCTGTTGGTGGGCAGGGATTAAGCAGGAATTTGGCATTCCCTACAATCCCCAAAGTCAGGAGTAGTAGAATCTATGAATAAAGAATTAAAGAAAATTATAGGACAGGTAGAGAACAAGCTGAACATCTTAAGACAGCAGTACAAATGGCAGTATTCATC

TAGAAGCAGAAGTTATTCCAGCAGAAACAGGACAGGAAACAGCATACTTTATCTTAAAACTAGCAGGAAGATGGCCAGTAAAAACAATACACACAGACAATGGCAGCAATTTTACCAGCAATACAGTTAAGGCCGCCTGTTGGTGGGCAGGGATTAATCAGGAATTTGGCATTCCCTACAATCCCCAAAGTCAGGAGTAGTAGAATCTATGAATAAAGAATTAAAGAAAATTATAGGACAGGTAGAGAACAAGCTGAACATCTTAAGACAGCAGTACAAATGGCAGTATTCATC

TAGAAGCAGAAGTTATTCCAGCAGAAACAGGACAGGAAACAGCATACTTTATCTTAAAACTAGCAGGAAGATGGCCAGTAAAAACAATACACACAGACAATGGCAGCAATTTTACCAGTAATACAGTTAAGGCCGCCTGTTGGTGGGCAGGGATTAAGCAGGAATTTGGCATTCCCTACAATCCCCAAAGTCAGGAGTAGTAGAATCTATGAATAAAGAATTAAAGAAAATTATAGGACAGGTAGAGAACAAGCTGAACATCTTAAGACAGCAGTACAAATGGCAGTATTCATC

TAGAAGCAGAAGTTATTCCAGCAGAAACAGGACAGGAAACAGCATACTTTATCTTAAAACTAGCAGGAAGATGGCCAGTAAAAACAATACACACAGACAATGGCAGCAATTTTACCAGTAATACAGTTAAGGCCGCCTGTTGGTGGGCAGGGATTAAGCAGGAATTTGGCATTCCCTACAATCCCCAAAGTCAGGAGTAGTAGAATCTATGAATAAAAAATTAAAGAAAATTATAGGACAGGTAGAGAACAAGCTGAACATCTTAAGACAGCAGTACAAATGGCAGTATTCATC

TAGAAGCAGAAGTTATTCCAGCAGAAACAGGACAGGAAACAGCATACTTTATCTTAAAACTAGCAGGAAGATGGCCAGTAAAAACAATACACACAGACAATGGCAGCAATTTTTCCAGTAATACAGTTAAGGCCGCCTGTTGGTGGGCAGGGATTAAGCAGGAATTTGGTATTCCCTACAATCCCCAAAGTCAGGAGTAGTAGAATCTATGAATAAAGAATTAAAGAAAATTATAGGACAGGTAGAGAACAAGCTGAACATCTTAAGACAGCAGTACAAATGGCAGTATTCATC

TAGAAGCAGAAGTTATTCCAGCAGAAACAGGACAGGAAACAGCATACTTTATCTTAAAACTAGCAGGAAGATGGCCAGTAAAAACAATACACACAGACAATGGCAGCAATTTTACCAGTAACACAGTTAAGGCCGCCTGTTGGTGGGCAGGGATTAATCAGGAATTTGGCATTCCCTACAATCCCCAAAGTCAGGAGTAGTAGAATCTATGAATAAAGAATTAAAGAAAATTATAGGACAGGTAGAGAACAAGCTGAACATCTTAAGACAGCAGTACAAATGGCAGTATTCATC

TAGAAGCAGAAGTTATTCCAGCAGAAACAGGACAGGAAACAGCATACTTTATCTTAAAACTAGCAGGAAGATGGCCAGTAAAAACAATACACACAGACAATGGCAGCAATTTTACCAGTAATACAGTTAAGGCCGCCTGTTGGTGGGCAGGGATTAAGCAGGAATTTGGCATTCCCTACAATCCCCAAAGTCAGGAGTAGTAGAATCTATGAATAAAGAATTAAAAAAAATTATAGGACAGGTAGAGAACAAGCTGAACATCTTAAGACAGCAGTACAAATGGCAGTATTCATC

TAGAAGCAGAAGTTATTCCAGCAGAAACAGGACAGGAAACAGCATACTTTATCTTAAAACTAGCAGGAAGATGGCCAGTAAAAACAATACACACAGACAATGGCAGCAATTTTACCAGTAATACAGTGAGGCCGCCTGTTGGTGGGCAGGGATTAATCAGGAATTTGGCATTCCCTACAATCCCCAAAGTCAGGAGTAGTAGAATCTATGAATAAAGAATTAAAGAAAATTATAGGACAGGTAGAGAACAAGCTGAACATCTTAAGACAGCAGTACAAATGGCAGTATTCATC

TAGAAGCAGAAGTTATTCCAGCAGAAACAGGACAGGAAACAGCATACTTTATCTTAAAACTAGCAGGAAGATGGCCAGTAAAAACAATACACACAGACAATGGCAGCAATTTTACCAGTAATACAGTTAAGGCCGCCCGTTGGTGGGCAGGGATTAAGCAGGAATTTGGCATTCCCTACAATCCCCAAAGTCAGGAGTAGTAGAATCTATGAATAAAGAATTAAAGAAAATTATAGGACAGGTAGAGAACAAGCTGAACATCTTAAGACAGCAGTACAAATGGCAGTATTCATC

TAGAAGCAGAAGTTATTCCAGCAGAAACAGGACAGGAAACAGCATACTTTATCTTAAAACTAGCAGGAAGATGGCCAGTAAAAACAATACACACAGACAATGGCAGCAATTTTACCAGTAATACAGTTAAGGCCGCCTGTTGGTGGGCAGGGATTAAGCAGGAATTTGGCATTCCCTACAATCCCCAAAGTCAGGAGTAATAGAATCTATGAATAAAGAATTAAAGAAAATTATAGGACAGGTAGAGAACAAGCTGAACATCTTAAGACAGCAGTACAAATGGCAGTATTCATC

TAGAAGCAGAAGTTATTCCAGCAGAAACAGGACAGGAAACAGCATACTTTATCTTAAAACTAGCAGGAAGATGGCCAGTAAAAACAATACACACAGACAATGGCAGCAATTTTACCAGTAATACAGTTAAGGCCGCCTGTTGGTGGGCAGGGATTAAGCAGGAATTTGGCATTCCCTACAATCCCCAAAGTCAGGAGTAGTAGAATCTATGAACAAAGAATTAAAGAAAATTATAGGACAGGTAGAGAACAAGCTGAACATCTTAAGACAGCAGTACAAATGGCAGTATTCATC

TAGAAGCAGAAGTTATTCCAGCAGAAACAGGACAGGAAACAGCATACTTTATCTTAAAACTAGCAGGAAGATGGCCAGTAAAAACAATATACACAGACAATGGCAGCAATTTTACCAGTAATACAGTTAAGGCCGCCTGTTGGTGGGCAGGGATTAAGCAGGAATTTGGCATTCCCTACAATCCCCAAAGTCAGGAGTAGTAGAATCTATGAATAAAGAATTAAAGAAAATTATAGGACAGGTAGAGAACAAGCTGAACATCTTAAGACAGCAGTACAAATGGCAGTATTCATC

TAGAAGCAGAAGTTATTCCAGCAGAAACAGGACAGGAAACAGCATACTTTATCTTAAAACTAGCAGGAAGATGGCCAGTAAAAACAATACACACAGACAATGGCAGCAATTTTACCAGTAATACAGTTAAGGCCGCCTGCTGGTGGGCAGGGATTAAGCAGGAATTTGGCATTCCCTACAATCCCCAAAGTCAGGAGTAGTAGAATCTATGAATAAAGAATTAAAGAAAATTATAGGACAGGTAGAGAACAAGCTGAACATCTTAAGACAGCAGTACAAATGGCAGTATTCATC

TAGAAGCAGAAGTTATTCCAGCAGAAACAGGACAGGAAACAGCATACTTTATCTTAAAACTAGCAGGAAGATGGCCAGTAAAAACAATACACACAGACAATGGCAGCAATTTTACCAGTAATACAGTTAAGGCCGCCTGTTGGTGGGCAGGGATTAAGCAGGAATTTGGCATTCCCTACAATCCCCAAAGTCAGGAGTAGTAGAATCTATGAATAAAGAATTAAAGAAAATTATAGGACAGGTAGAGAACAAGCTGAACATCTTAAGACAGCAGTACAAATGGCAGTATTCATC

TAGAAGCAGAAGTTATTCCAGCAGAAACAGGACAGGAAACAGCATACTTTATCTTAAAACTAGCAGGAAGATGGCCAGTAAAAACAATACATACAGACAATGGCAGCAATTTTACCAGTAATACAGTTAAGGCCGCCTGTTGGTGGGCAGGGATTAATCAGGAATTTGGCATTCCCTACAATCCCCAAAGTCAGGAGTAGTAGAATCTATGAATAAAGAATTAAAGAAAATTATAGGACAGGTAGAGAACAAGCTGAACATCTTAAGACAGCAGTACAAATGGCAGTATTCATC

TAGAAGCAGAAGTTATTCCAGCAGAAACAGGACAGGAAACAGCATACTTTATCTTAAAACTAGCAGGAAGATGGCCAGTAAAAACAATACACACAGACAATGGCAGCAATTTTACCAGTAATACAGTGAAGGCCGCCTGTTGGTGGGCAGGGATTAAGCAGGAATTTGGCATTCCCTACAATCCCCAAAGTCAGGAGTAGTAGAATCTATGAATAAAGAATTAAAGAAAATTATAGGACAGGTAGAGAACAAGCTGAACATCTTAAGACAGCAGTACAAATGGCAGTATTCATC

TAGAAGCAGAAGTTATTCCAGCAGAAACAGGACAGGAAACAGCATACTTTATCTTAAAACTAGCAGGAAGATGGCCAGTAAAAACAATACACACAGATAATGGCGGCAATTTTACCAGTAATACAGTTAAGGCCGCCTGTTGGTGGGCAGGGATTAAGCAGGAATTTGGCATTCCCTACAATCCCCAAAGTCAGGAGTAGTAGAATCTATGAATAAAGAATTAAAGAAAATTATAGGACAGGTAGAGAACAAGCTGAACATCTTAAGACAGCAGTACAAATGGCAGTATTCATC

TAGAAGCAGAAGTTATTCCAGCAGAAACAGGACAGGAAACAGCATACTTTATCTTAAAATTAGCAGGAAGATGGCCAGTAAAAACAATACACACAGACAATGGCAGCAATTTTACCAGTAATACAGTTAAGGCCGCCTGTTGGTGGGCAGGGATTAATCAGGAATTTGGCATTCCCTACAATCCCCAAAGTCAGGAGTAGTAGAATCTATGAATAAAGAATTAAAGAAAATTATAGGACAGGTAGAGAACAAGCTGAACATCTTAAGACAGCAGTACAAATGGCAGTATTCATC

TAGAAGCAGAAGTTATTCCAGCAGAAACAGGACAGGAAACAGCATACTTTATCTTAAAACTAGCAGGAAGATGGCCAGTAAAAACAATACACACAGACAATGGCAGCAATTTTACCAGTAATACAGTTAAGGCCGCCTGTTGGTGGGCAGGGATTAGGCAGGAATTTGGCATTCCCTACAATCCCCAAAGTCAGGAGTAGTAGAATCTATGAATAAAGAATTAAAGAAAATTATAGGACAGGTAGAGAACAAGCTGAACATCTTAAGACAGCAGTACAAATGGCAGTATTCATC

TAGAAGCAGAAGTTATTCCAGCAGAAACAGGACAGGAAACAGCATACTTTATCTTAAAACTAGCAGGAAGATGGCCAGTAAAAACAATACACACAGACAATGGCAGTAATTTTACCAGTAATACAGTTAAGGCCGCCTGTTGGTGGGCAGGGATTAATCAGGAATTTGGCATTCCCTACAATCCCCAAAGTCAGGAGTAGTAGAATCTATGAATAAAGAATTAAAGAAAATTATAGGACAGGTAGAGAACAAGCTGAACATCTTAAGACAGCAGTACAAATGGCAGTATTCATC

TAGAAGCAGAAGTTATTCCAGCAGAAACAGGACAGGAAACAGCATACTTTATCTTAAAACTAGCAGGAAGATGGCCAGTAAAAACAATACACACAGACAATGGCAGCAATTTTACCAGTAATACAGTTAAGGCCGCCTGTTGGTGGGCAGGGATTAAGCAGGAATTTGGCATTCCCTACAATCCCCAAAGTCAGGAGTAGTAGAATCTATGAATAAAGAGTTAAAGAAAATTATAGGACAGGTAGAGAACAAGCTGAACATCTTAAGACAGCAGTACAAATGGCAGTATTCATC

TAGAAGCAGAAGTTATTCCAGCAGAAACAGGACAGGAAACAGCATACTTTATCTTAAAACTAGCAGGAAGATGGCCAGTGAAAACAATACACACAGACAATGGCAGCAATTTTACCAGTAATACAGTTAAGGCCGCCTGTTGGTGGGCAGGGATTAAGCAGGAATTTGGCATTCCCTACAATCCCCAAAGTCAGGAGTAGTAGAATCTATGAATAAAGAATTAAAGAAAATTATAGGACAGGTAGAGAACAAGCTGAACATCTTAAGACAGCAGTACAAATGGCAGTATTCATC

TAGAAGCAGAAGTTATTCCAGCAGAAACAGGACAGGAAACAGCATACTTTATCTTAAAACTAGCAGGAAGATGGCCAGTAAAAACAATACACACAGACAATGGCAGCAATTTTACCAGTAATACAGTTAAGGCCGCCTGTTGGTGGGCAGGGATTAAGCAGGAATTTGGCATTCCCTACAATCCCCAAAGTCAGGAGTGGTAGAATCTATGAATAAAGAATTAAAGAAAATTATAGGACAGGTAGAGAACAAGCTGAACATCTTAAGACAGCAGTACAAATGGCAGTATTCATC

TAGAAGCAGAAGTTATTCCAGCAGAGACAGGACAGGAAACAGCATACTTTATCTTAAAACTAGCAGGAAGATGGCCAGTAAAAACAATACACACAGACAATGGCAGCAATTTTACCAGTAATACAGTTAAGGCCGCCTGTTGGTGGGCAGGGATTAAGCAGGAATTTGGCATTCCCTACAATCCCCAAAGTCAGGAGTAGTAGAATCTATGAATAAAGAATTAAAGAAAATTATAGGACAGGTAGAGAACAAGCTGAACATCTTAAGACAGCAGTACAAATGGCAGTATTCATC

TAGAAGCAGAAGTTATTCCAGCAGAAACAGGACAGGAAACAGCATACTTTATATTAAAACTAGCAGGAAGATGGCCAGTAAAAACAATACACACAGACAATGGCAGCAATTTTACCAGTAATACAGTTAAGGCCGCCTGTTGGTGGGCAGGGATTAAGCAGGAATTTGGCATTCCCTACAATCCCCAAAGTCAGGAGTAGTAGAATCTATGAATAAAGAATTAAAGAAAATTATAGGACAGGTAGAGAACAAGCTGAACATCTTAAGACAGCAGTACAAATGGCAGTATTCATC

TAGAAGCAGAAGTTATTCCAGCAGAAACAGGACAGGAAACAGCATACTTTATCTTAAAACTAGCAGGAAGATGGCCAGTAAAAACAATACACACAGACAATGGCAGCAATTTTACCAGTAATACAGTTAAGGCCGCCTGTTGGTGGGCAGGGATTAAGCAGGAATTTGGCATTCCCTACAATCCCCAAAGTCAGGAGTAGTAGAATCTATAAATAAAGAATTAAAGAAAATTATAGGACAGGTAGAGAACAAGCTGAACATCTTAAGACAGCAGTACAAATGGCAGTATTCATC

TAGAAGCAGAAGTTATTCCAGCAGAAACAGGACAGGAAACAGCATACTTTATCTTAAAACTAGCAGGAAGATGGCCAGTAAGAACAATACACACAGACAATGGCAGCAATTTTACCAGTAATACAGTTAAGGCCGCCTGTTGGTGGGCAGGGATTAAGCAGGAATTTGGCATTCCCTACAATCCCCAAAGTCAGGAGTAGTAGAATCTATGAATAAAGAATTAAAGAAAATTATAGGACAGGTAGAGAACAAGCTGAACATCTTAAGACAGCAGTACAAATGGCAGTATTCATC

TAGAAGCAGAAGTTATTCCAGCAGAAACAGGACAGGAAACAGCATACTTTATATTAAAACTAGCAGGAAGATGGCCAGTAAAAACAATACACACAGACAATGGCAGCAATTTTACCAGTAATACAGTTAAGGCCGCCTGTTGGTGGGCAGGGATTAAGCAGGAATTTGGCATTCCCTACAATCCCCAAAGTCAGGAGTAGTAGAATCTATGAATAAAGAATTAAAGAAAATTATAGGACAGGTAGAGAACAAGCTGAACATCTTAAGACAGCAGTACAAATGGCAGTATTCATC

TAGAAGCAGAAGTTATTCCAGCAGAAACAGGACAGGAAACAGCATACTTTATCTTAAAACTAGCAGGAAGATGGCCAGTAAAAACAATACACACAGACAATGGCAGCAATTTTACTAGTAACACAGTTAAGGCCGCCTGTTGGTGGGCAGGGATTAATCAGGAATTTGGCATTCCCTACAATCCCCAAAGTCAGGAGTAGTAGAATCTATGAATAAAGAATTAAAGAAAATTATAGGACAGGTAGAGAACAAGCTGAACATCTTAAGACAGCAGTACAAATGGCAGTATTCATC

TAGAAGCAGAAGTTATTCCAGCAGAAACAGGACAGGAAACAGCATACTTTATCTTAAAACTAGCAGGAAGATGGCCCGTAAAAACAATACACACAGACAATGGCAGCAATTTTACCAGTAATACAGTTAAGGCCGCCTGTTGGTGGGCAGGGATTAAGCAGGAATTTGGCATTCCCTACAATCCCCAAAGTCAGGAGTAGTAGAATCTATGAATAAAGAATTAAAGAAAATTATAGGACAGGTAGAGAACAAGCTGAACATCTTAAGACAGCAGTACAAATGGCAGTATTCATC

TAGAAGCAGAAGTTATTCCAGCAGAAACAGGACAGGAAGAAACAGCATACTTTATCTTAAAACTAGCAGGAAGATGGCCAGTAAAAACAATACACACAGACAATGGCGGCAATTTTACCAGTAATACAGTTAAGGCCGCCTGTTGGTGGGCAGGGATTAAGCAGGAATTTGGCATTCCCTACAATCCCCAAAGTCAGGAGTAGTAGAATCTATGAATAAAGAATTAAAGAAAATTATAGGACAGGTAGAGAACAAGCTGAACATCTTAAGACAGCAGTACAAATGGCAGTATTCATC

TAGAAGCAGAAGTTATTCCAGCAGAAACAGGACAGGAAACAGCATACTTTATCTTAAAACTAGCAGGAAGATGGCCAGTAAAAACAATACACACAGACAATGGCAGCAATTTTACCAGTAATACAGTTAAGGCCGCCTGTTGGTGGGCAGGGATTAAGCAGGAATTTGGCATTCCCTACAATCCCCAAAGGCAGGAGTAGTAGAATCTATGAATAAAGAATTAAAGAAAATTATAGGACAGGTAGAGAACAAGCTGAACATCTTAAGACAGCAGTACAAATGGCAGTATTCATC

TAGAAGCAGAAGTTATTCCAGCAGAAACAGGACAGGAAACAGCATACTTTATCTTAAAACTAGCAGGAAGATGGCCAGTAAAAACAATACACACAGACAATGGCAGCAATTTTACCAGTAATACAGTTAAGGCCGCCTGTTGGTGGGCAGGGATTAAGCAGAATTTGGCATTCCCTACAATCCCCAAAGTCAGGAGTAGTAGAATCTATGAATAAAGAATTAAAGAAAATTATAGGACAGGTAGAGAACAAGCTGAACATCTTAAGACAGCAGTACAAATGGCAGTATTCATC

TAGAAGCAGAAGTTATTCCAGCAGAAACAGGACAGGAAACAGCATACTTTATCTTAAAACTAGCAGGAAGATGGCCAGTAAAAACAATACACACAGACAATGGCAGCAATTTTACCAGTAATACAGTTAAGGCCGCCTGTTGGTGGGCAGGGATTAAGCAGGAATTTGGCATTCCCTACAATCCCCAAAGTCAGGAGTAGTAGAATCTATGAATAAAGAATTAAAGAAAATTATAGGACAGGTAGAGAACAAGCTGAACACCTTAAGACAGCAGTACAAATGGCAGTATTCATC

TAGAAGCAGAAGTTATTCCAGCAGAAACAGGACAGGAAACAGCATACTTTATCTTAAAACTAGCAGGAAGATGGCCAGTAAAAACAATACACACAGACAATGGCAGCAATTTTACCAGTAATACAGTTAAGGCCGCCTGTTGGTGGGCAGGGATTAAGCAGGAATTTGGCATTCCCTACAATCCCCAAAGTCAGGAGTAGTAGAATCTATGAATAAAGAATTAAAGAAAATTATAGGACAGGTAGAGAACAAGCTGAACATCTTAAGACAGCAGTACAAATGGCAGTATTCATC

TAGAAGCAGAAGTTATTCCAGCAGAAACAGGACAGGAAACAGCATACTTTATCTTAAAACTAGCAGGAAGATGGCCAGTAAAAACAATACACACAGACAATGGCAGCAATTTTACCAGTAATACGGTTAAGGCCGCCTGTTGGTGGGCAGGGATTAATCAGGAATTTGGCATTCCCTACAATCCCCAAAGTCAGGAGTAGTAGAATCTATGAATAAAGAATTAAAGAAAATTATAGGACAGGTAGAGAACAAGCTGAACATCTTAAGACAGCAGTACAAATGGCAGTATTCATC

TAGAAGCAGAAGTTATTCCAGCAGAAACAGGACAGGAAACAGCATACTTTATCTTAAAACTAGCAGGAAGATGGCCAGTAAAAACAATACACACAGACAATGGCAGCAATTTTACCAGCAATACAGTTAAGGCCGCCTGTTGGTGGGCAGGGATTAAACAGGAATTTGGCATTCCCTACAATCCCCAAAGTCAGGAGTAGTAGAATCTATGAATAAAGAATTAAAGAAAATTATAGGACAGGTAGAGAACAAGCTGAACATCTTAAGACAGCAGTACAAATGGCAGTATTCATC

TAGAAGCAGAAGTTATTCCAGCAGAAACAGGACAGGAAACAGCATACTTTATCTTAAAACTAGCAGGAAGATGGCCAGTAAAAACAATACACACAGACAATGGCAGCAATTTTACCAGTAATACAGTAAAGGCCGCCTGTTGGTGGGCAGGGATTAAGCAGGAATTTGGCATTCCCTACAATCCCCAAAGTCAGGAGTAGTAGAATCTATGAATAAAGAATTAAAGAAAATTATAGGACAGGTAGAGAACAAGCTGAACATCTTAAGACAGCAGTACAAATGGCAGTATTCATC

TAGAAGCAGAAGTTATTCCAGCAGAAACAGGACAGGAAACAGCATACTTTATATTAAAACTAGCAGGAAGATGGCCAGTAAAAACAATACACACAGACAATGGCAGCAATTTTACCAGTAATACAGTTAAGGCCGCCTGTTGGTGGGCAGGGATTAAGCAGGAATTTGGCATTCCCTACAATCCCCAAAGTCAGGAGTAGTAGAATCTATAAATAAAGAATTAAAGAAAATTATAGGACAGGTAGAGAACAAGCTGAACATCTTAAGACAGCAGTACAAATGGCAGTATTCATC

TAGAAGCAGAAGTTATTCCAGCAGAAACAGGACAGGAAACAGCATACTTTATCTTAAAACTAGCAGGAAGATGGCCAGTAAAAACAATACACACAGACAATGGCAGCAATTTTACCAGTAATACAGTTAAGGCCGCCTGTTGGTGGGCAGGGATTAAGCAGGAATTTGGCATTCCCTACAATCCCCAAAGTCAGGAGTAGTAGAATCTATGAATAAAGAATTAAAGAAAATTATAGGACAGGTAGAGAACAAGCTGAACATCTTAAGACAGCAGTACAAATGGCAGTATT-ATC

TAGAAGCAGAAGTTATTCCAGCAGAAACAGGACAGGAAACAGCATACTTTATCTTAAAACTAGCAGGAAGATGGCCAGTAAAAACAATACACACAGACAATGGCAGCAATTTTACCAGTAATACAGTTAAGGCCGCCTGTTGGTGGGCAGGGATTAATCAGGAATTTGGCATTCCCTACAATCCCCAAAGTCAGGAGTAGTAGAATCTATGAATAAAAAATTAAAGAAAATTATAGGACAGGTAGAGAACAAGCTGAACATCTTAAGACAGCAGTACAAATGGCAGTATTCATC

TAGAAGCAGAAGTTATTCCAGCAGAAACAGGACAGGAAACAGCATACTTTATCTTAAAACTAGCAGGAAGATGGCCAGTAAAAACAATACACACAGACAATGGCAGCAATTTTACCAGTAATACAGTTAAGGCCGCCTGTTGGTGGGCAGGGATTAAGCAGGAATTTGGCATTCCCTACAATCCCCAAAGTCAGGAGTAGTAGAATCTATGAATAAAGAATTAAAGAAAATTATAGGACAGGTAAAGAACAAGCTGAACATCTTAAGACAGCAGTACAAATGGCAGTATTCATC

TAGAAGCAGAAGTTATTCCAGCAGAAACAGGACAGGAAACAGCCTACTTTATCTTAAAACTAGCAGGAAGATGGCCAGTAAAAACAATACACACAGACAATGGCAGCAATTTTACCAGTAATACAGTTAAGGCCGCCTGTTGGTGGGCAGGGATTAAGCAGGAATTTGGCATTCCCTACAATCCCCAAAGTCAGGAGTAGTAGAATCTATGAATAAAGAATTAAAGAAAATTATAGGACAGGTAGAGAACAAGCTGAACATCTTAAGACAGCAGTACAAATGGCAGTATTCATC

TAGAAGCAGAAGTTATTCCAGCAGAAACAGGACAGGAAACAGCATACTTTATCTTAAAACTAGCAGGAAGATGGCCAGTAAAAACAATACACACAGACAATGGCAGCAATTTTACCAGTAATACAGTTAAGGCCGCCTGTTGGTGGGCAGGGATTAATCAGGAATTTGGCATTCCCTACAATCCCC-AAGTCAGGAGTAGTAGAATCTATGAATAAAGAATTAAAGAAAATTATAGGACAGGTAGAGAACAAGCTGAACATCTTAAGACAGCAGTACAAATGGCAGTATTCATC

TAGAAGCAGAAGTTATTCCAGCAGAGACAGGACAGGAAACAGCATACTTTATCTTAAAACTAGCAGGAAGATGGCCAGTAAAAACAATACATACAGACAATGGCAGCAATTTTACCAGTAATACAGTTAAGGCCGCCTGTTGGTGGGCAGGGATTAAGCAGGAATTTGGCATTCCCTACAATCCCCAAAGTCAGGAGTAGTAGAATCTATGAATAAAGAATTAAAGAAAATTATAGGACAGGTAGAGAACAAGCTGAACATCTTAAGACAGCAGTACAAATGGCAGTATTCATC

TAGAAGCAGAAGTTATTCCAGCAGAAACAGGACAGGAAACAGCATACTTTATCTTAAAACTAGCAGGAAGATGGCCAGT--AAACAATACACACAGACAATGGCAGCAATTTTACCAGCAATACAGTTAAGGCCGCCTGTTGGTGGGCAGGGATTAAGCAGGAATTTGGCATTCCCTACAATCCCCAAAGTCAGGAGTAGTAGAATCTATGAATAAAGAATTAAAGAAAATTATAGGACAGGTAGAGAACAAGCTGAACATCTTAAGACAGCAGTACAAATGGCAGTATTCATC

TAGAAGCAGAAGTTATTCCAGCAGAAACAGGACAGGAAACAGCATACTTTATCTTAAAACTAGCAGGAAGATGGCCAGTAAAAACAATACACACAGACAATGGCAGCAATTTCACCAGTAATACAGTTAAGGCCGCCTGTTGGTGGGCAGGGATTAATCAGGAATTTGGCATTCCCTACAATCCCCAAAGTCAGGAGTAGTAGAATCTATGAATAAAGAATTAAAGAAAATTATAGGACAGGTAGAGAACAAGCTGAACATCTTAAGACAGCAGTACAAATGGCAGTATTCATC

TAGAAGCAGAAGTTATTCCAGCAGAAACAGGACAGGAAACAGCATACTTTATCTTAAAACTAGCAGGAAGATGGCCAGTAAAAACAATACACACAGACAATGGCAGCAATTTTACTAGTAATACAGTTAAGGCCGCCTGTTGGTGGGCAGGGATTAAGCAGGAATTTGGCATTCCCTACAATCCCCAAAGTCAGGAGTAGTAGAATCTATGAATAAAGAATTAAAGAAAATTATAGGACAGGTAGAGAACAAGCTGAACATCTTAAGACAGCAGTACAAATGGCAGTATTCATC

TAGAAGCAGAAGTTATTCCAGCAGAAACAGGACAGGAAACAGCATACTTTATCTTAAAACTAGCAGGAAGATGGCCAGTAAAAACAATATACACAGACAATGGCAGCAATTTTACCAGTAATACAGTTAAGGCCGCCTGTTGGTGGGCAGGGATTAAGCAGGAATTTGGCATTCCCTACAATCCCCAAAGTCAGGAGTAGTAGAATCTATGAATAAAGAATTAAAGAAAATTATAGGACAGGTAGAGAACAAGCTGAACATCTTAAGACAGCAGTACAAATGGCAGTATTCATC

TAGAAGCAGAAGTTATTCCAGCAGAAACAGGACAGGAAACAGCATACTTCATCTTAAAACTAGCAGGAAGATGGCCAGTAAAAACAATACACACAGACAATGGCAGCAATTTTACCAGTAATACAGTTAAGGCCGCCTGTTGGTGGGCAGGGATTAAGCAGGAATTTGGCATTCCCTACAATCCCCAAAGTCAGGAGTAGTAGAATCTATGAATAAAGAATTAAAGAAAATTATAGGACAGGTAGAGAACAAGCTGAACATCTTAAGACAGCAGTACAAATGGCAGTATTCATC

TAGAAGCAGAAGTTATTCCAGCAGAAACAGGACAGGAAACAGCATACTTTATCtTAAAACTAGCAGGAAGATGGCCAGTAAAAACAATACACACAGACAATGGCAGCAATTTTACCAGTAATACAGTTAAGGCCGCCTGTTGGTGGGCAGGGATTAAGCAGGAATTTGGCATTCCCTACAATCCCCAAAGTCAGGAGTAGTAGAATCTATGAATAAAGAATTAAAGAAAATTATAGGACAGGTAGAGAACAAGCTGAACATCTTAAGACAGCAGTACAAATGGCAGTATTCATC

TAGAAGCAGAAGTTATTCCAGCAGAAACAGGACAGGAAACAGCATACTTTATCCTAAAACTAGCAGGAAGATGGCCAGTAAAAACAATACACACAGACAATGGCAGCAATTTTACCAGTAATACAGTTAAGGCCGCCTGTTGGTGGGCAGGGATTAAGCAGGAATTTGGCATTCCCTACAATCCCCAAAGTCAGGAGTAGTAGAATCTATGAATAAAGAATTAAAGAAAATTATAGGACAGGTAGAGAACAAGCTGAACATCTTAAGACAGCAGTACAAATGGCAGTATTCATC

TAGAAGCAGAAGTTATTCCAGCAGAGACAGGACAGGAAACAGCATACTTTATCTTAAAACTAGCAGGAAGATGGCCAGTAAAAACAATACATACAGACAATGGCAGCAATTTTACCAGTAATACAGTTAAGGCCGCCTGTTGGTGGGCAGGGATTAAGCAGGAATTTGGCATTCCCTACAATCCCCAAAGTCAGGAGTAGTAGAATCTATGAATAAAGAATTAAAGAAAATTATAGGACAGGTAGAGAACAAGCTGAACATCTTAAGACAGCAGTACAAATGGCAGTATTCATC

TAGAAGCAGAAGTTATTCCAGCAGAAACAGGACAGGAAACAGCATACTTTATCTTAAAACTAGCAGGAAGATGGCCAGTAAAAACAATACACACAGACAATGGCAGCAATTTTACCAGTAATACAGTTAAGGCCGCCTGTTGGTGGGCAGGGATTAATCAGGAATTTGGCATTCCCTACAATCCCCAAAGTCAGGAGTAGTAGAATCTATGAATAAAGAATTAAAAAAAATTATAGGACAGGTAGAGAACAAGCTGAACATCTTAAGACAGCAGTACAAATGGCAGTATTCATC

TAGAAGCAGAAGTTATTCCAGCAGAAACAGGACAGGAAACAGCATATTTTATCTTAAAACTAGCAGGAAGATGGCCAGTAAAAACAATACACACAGACAATGGCAGCAATTTTACCAGTAATACAGTTAAGGCCGCCTGTTGGTGGGCAGGGATTAAACAGGAATTTGGCATTCCCTACAATCCCCAAAGTCAGGAGTAGTAGAATCTATGAATAAAGAATTAAAGAAAATTATAGGACAGGTAGAGAACAAGCTGAACATCTTAAGACAGCAGTACAAATGGCAGTATTCATC

TAGAAGCAGAAGTTATTCCAGCAGAAACAGGACAGGAAACAGCATACTTTATCCTAAAACTAGCAGGAAGATGGCCAGTAAAAACAATACACACAGACAATGGCAGCAATTTTACCAGTAATACAGTTAAGGCCGCCTGTTGGTGGGCAGGGATTAATCAGGAATTTGGCATTCCCTACAATCCCCAAAGTCAGGAGTAGTAGAATCTATGAATAAAGAATTAAAGAAAATTATAGGACAGGTAGAGAACAAGCTGAACATCTTAAGACAGCAGTACAAATGGCAGTATTCATC

| Reads | AvgQScore | Type | Pct | IndelLength |
|---|---|---|---|---|
| 6183 | 36 | 19 Base Changes | 38.35 | 0 |
| 2679 | 36 | 20 Base Changes | 16.61 | 0 |
| 496 | 36 | 20 Base Changes | 3.08 | 0 |
| 480 | 36 | 20 Base Changes | 2.98 | 0 |
| 224 | 36 | 21 Base Changes | 1.39 | 0 |
| 191 | 36 | 18 Base Changes | 1.18 | 0 |
| 189 | 36 | 20 Base Changes | 1.17 | 0 |
| 169 | 36 | 20 Base Changes | 1.05 | 0 |
| 154 | 36 | 20 Base Changes | 0.96 | 0 |
| 126 | 36 | 21 Base Changes | 0.78 | 0 |
| 103 | 36 | 20 Base Changes | 0.64 | 0 |
| 102 | 36 | 20 Base Changes | 0.63 | 0 |
| 86 | 36 | 19 Base Changes | 0.53 | 0 |
| 82 | 36 | 18 Base Changes | 0.51 | 0 |
| 77 | 36 | 20 Base Changes | 0.48 | 0 |
| 74 | 36 | 20 Base Changes | 0.46 | 0 |
| 61 | 36 | 20 Base Changes | 0.38 | 0 |
| 60 | 36 | 21 Base Changes | 0.37 | 0 |
| 59 | 36 | 20 Base Changes | 0.37 | 0 |
| 57 | 36 | 18 Base Changes | 0.35 | 0 |
| 54 | 36 | 21 Base Changes | 0.33 | 0 |
| 53 | 36 | 21 Base Changes | 0.33 | 0 |
| 53 | 36 | 20 Base Changes | 0.33 | 0 |
| 52 | 36 | 18 Base Changes | 0.32 | 0 |
| 49 | 37 | 21 Base Changes | 0.3 | 0 |
| 49 | 36 | 20 Base Changes | 0.3 | 0 |
| 45 | 36 | 21 Base Changes | 0.28 | 0 |
| 45 | 37 | 20 Base Changes | 0.28 | 0 |
| 44 | 36 | 20 Base Changes | 0.27 | 0 |
| 44 | 36 | 20 Base Changes | 0.27 | 0 |
| 43 | 36 | 21 Base Changes | 0.27 | 0 |
| 42 | 36 | 20 Base Changes | 0.26 | 0 |
| 37 | 35 | 20 Base Changes | 0.23 | 0 |
| 37 | 36 | 19 Base Changes | 0.23 | 0 |
| 36 | 36 | 21 Base Changes | 0.22 | 0 |
| 35 | 36 | 21 Base Changes | 0.22 | 0 |
| 35 | 36 | 19 Base Changes | 0.22 | 0 |
| 34 | 36 | 20 Base Changes | 0.21 | 0 |
| 34 | 36 | 21 Base Changes | 0.21 | 0 |
| 33 | 36 | 20 Base Changes | 0.2 | 0 |
| 33 | 36 | 20 Base Changes | 0.2 | 0 |
| 32 | 36 | 20 Base Changes | 0.2 | 0 |
| 32 | 35 | 20 Base Changes | 0.2 | 0 |
| 31 | 36 | 19 Base Changes | 0.19 | 0 |
| 31 | 36 | 20 Base Changes | 0.19 | 0 |
| 29 | 36 | 20 Base Changes | 0.18 | 0 |
| 28 | 36 | 20 Base Changes | 0.17 | 0 |
| 28 | 36 | 22 Base Changes | 0.17 | 0 |
| 28 | 36 | 20 Base Changes | 0.17 | 0 |
| 27 | 36 | 21 Base Changes | 0.17 | 0 |
| 27 | 36 | 21 Base Changes | 0.17 | 0 |
| 26 | 36 | 20 Base Changes | 0.16 | 0 |
| 26 | 35 | 20 Base Changes | 0.16 | 0 |
| 26 | 36 | 18 Base Changes | 0.16 | 0 |
| 25 | 36 | 17 Base Changes | 0.16 | 0 |
| 25 | 36 | 21 Base Changes | 0.16 | 0 |
| 24 | 36 | 20 Base Changes | 0.15 | 0 |
| 24 | 35 | 20 Base Changes | 0.15 | 0 |
| 24 | 36 | Deletion | 0.15 | -1 |
| 24 | 36 | 21 Base Changes | 0.15 | 0 |
| 24 | 37 | 21 Base Changes | 0.15 | 0 |
| 24 | 36 | 20 Base Changes | 0.15 | 0 |
| 23 | 36 | Deletion | 0.14 | -1 |
| 23 | 36 | 19 Base Changes | 0.14 | 0 |
| 22 | 36 | Deletion | 0.14 | -2 |
| 22 | 36 | 19 Base Changes | 0.14 | 0 |
| 22 | 36 | 20 Base Changes | 0.14 | 0 |
| 22 | 36 | 20 Base Changes | 0.14 | 0 |
| 20 | 36 | 20 Base Changes | 0.12 | 0 |
| 20 | 36 | Insertion | 0.12 | 1 |
| 19 | 36 | 20 Base Changes | 0.12 | 0 |
| 19 | 36 | 21 Base Changes | 0.12 | 0 |
| 18 | 36 | 21 Base Changes | 0.11 | 0 |
| 18 | 36 | 18 Base Changes | 0.11 | 0 |
| 16 | 36 | 21 Base Changes | 0.1 | 0 |
